# Structural basis of the mechanism and inhibition of a human ceramide synthase

**DOI:** 10.1101/2023.12.02.569723

**Authors:** Tomas C. Pascoa, Ashley C.W. Pike, Christofer S. Tautermann, Gamma Chi, Michael Traub, Andrew Quigley, Rod Chalk, Saša Štefanic, Sven Thamm, Alexander Pautsch, Elisabeth P. Carpenter, Gisela Schnapp, David B. Sauer

## Abstract

Ceramides are bioactive sphingolipids that play pivotal roles in regulating cellular metabolism. Ceramides and dihydroceramides are synthesized by a family of six ceramide synthase enzymes (CerS), each with distinct specificity for the acyl-CoA substrate. Importantly, the acyl chain length plays a key role in determining the physiological function of ceramides, as well as their role in metabolic disease. Ceramide with an acyl chain length of 16 carbons (C16 ceramide) has been implicated in obesity, insulin resistance and liver disease, and the C16 ceramide-synthesizing CerS6 is regarded as an attractive drug target for obesity-associated disease. Despite their importance, the molecular mechanism underlying ceramide synthesis by CerS enzymes remains poorly understood. Here, we report cryo-electron microscopy structures of human CerS6, capturing covalent intermediate and product-bound states. These structures, together with biochemical characterization using intact protein and small molecule mass spectrometry, reveal that CerS catalysis proceeds via a ping-pong reaction mechanism involving a covalent acyl-enzyme intermediate. Notably, the product-bound structure was obtained upon reaction with the mycotoxin fumonisin B_1_, providing new insights into its inhibition of CerS. These results provide a framework for understanding the mechanisms of CerS function, selectivity, and inhibition, and open new directions for future drug discovery targeting the ceramide and sphingolipid pathways.

## Introduction

Ceramides are the precursors for the synthesis of complex sphingolipids and are bioactive lipids with crucial roles in signaling. In particular, ceramides have been proposed to act as key metabolic sensors to promote fatty acid utilization and storage during excessive fatty acid availability^1^. However, abnormal ceramide accumulation is associated with metabolic dysfunction, and elevated levels of ceramides have been observed in obesity-related metabolic disorders such as diabetes, non-alcoholic fatty liver disease (NAFLD) and non-alcoholic steatohepatitis (NASH)^2–4^.

Ceramides are composed of a sphingosine (d18:1) long-chain base with an N-linked acyl chain, the length of which is critical to the lipids’ biological functions and roles in pathophysiology^5^. For example, C16:0 ceramide is the most common ceramide in adipose tissue and its levels are elevated in this tissue of obese humans^2^. In addition, insulin resistance is correlated with plasma C16:0 and C18:0 ceramides, subcutaneous adipose tissue C16:0 ceramides, and hepatic C16:0 and C18:0 ceramides^6–8^. Moreover, the levels of total ceramides and C16:0, C22:0 and C24:1 dihydroceramides were found to be elevated in the liver of insulin-resistant patients with NASH^3^.

In mammals, *de novo* synthesis of ceramides is preceded by synthesis of dihydroceramides through the N-acylation of the sphingoid long-chain base sphinganine (dihydrosphingosine; d18:0) by one of six ceramide synthases (CerS1-6)^9^. The resulting dihydroceramides are subsequently converted into ceramides by one of two dihydroceramide Δ4-desaturases (DES1-2)^10^. Alternatively, CerS can directly re-acylate recycled sphingosine, in the salvage pathway^11^. Of these enzymes, recent observations from human and mouse studies have highlighted CerS6 as an attractive target for treating obesity-associated metabolic disease, including type 2 diabetes and NASH^2,5,12–14^. Deletion of *CerS6* in mice granted protection against diet-induced obesity, steatohepatitis, and insulin resistance^2,14^, and liver-specific deletion improved glucose tolerance and mitochondrial morphology^14^. Strikingly, this effect was shown to only occur in CerS6^Δ/Δ^ mice but not in CerS5^Δ/Δ^ mice, though both show a preference for C16-CoA and therefore primarily produce C16 ceramides^15^ and hepatic C16:0 ceramide levels were reduced in both knock-outs^14^. This surprising effect was found to be a result of the specific interaction of CerS6-derived C16:0 sphingolipids in mitochondria with the mitochondrial fission factor (Mff)^14^. This revealed that the subcellular localization of ceramide production can lead to drastically different physiological outcomes^14^. In further support of the therapeutic potential of inhibiting CerS6, ablation of the protein’s expression in an obese insulin resistance mouse model led to reduced body fat and improved oral glucose tolerance and insulin sensitivity^12^.

Central to its enzymatic function, CerS contain a TRAM-Lag1-CLN8 (TLC) homology domain^16^, which includes a 52-residue Lag1p motif containing conserved histidines and aspartates required for activity^17,18^. In addition, CerS2-6 contain a non-essential Hox-like domain between TM1-2^19^, flanked by two essential and conserved positively charged residues^20^. The activity of each CerS is subject to complex regulation, which includes phosphorylation^21^, glycosylation^22^, dimerisation^23^ and other protein-protein interactions^24,25^. The six mammalian CerS also have different acyl chain-length specificities and tissue expression patterns, further influencing the tissue-specific distribution of the different ceramides^5,9,26^. However, the molecular mechanisms underlying synthesis and regulation of ceramides and dihydroceramides remain unclear.

Despite the interest in pharmacologically reducing CerS6 activity, at present, no specific inhibitors have been described for this family member. The best characterized CerS inhibitor is fumonisin B_1_ (FB_1_), the most prevalent member of the fumonisin family of mycotoxins produced by *Fusarium* species. FB_1_ is a potential carcinogen and teratogen and therefore of significant concern as it is a common contaminant in maize, rice, other cereals, and cereal-based food stocks^27–29^. FB_1_ is a potent non-selective inhibitor of all human CerS, and is competitive towards both sphinganine and acyl-CoA substrates^30^. In addition, FTY720 (fingolimod, Gilenya), a prodrug administered to functionally antagonize the sphingosine-1-phosphate receptor 1 in the treatment of multiple sclerosis, is also a non-selective CerS inhibitor^31,32^. Further, a non-phosphorylatable analogue of FTY720, P053, has recently been shown to potently and selectively inhibit CerS1^33^, showcasing the possibility of achieving isoform-specific CerS inhibition.

Here, to understand the molecular basis of ceramide and dihydroceramide synthesis by the ceramide synthases, we employed a combination of cryogenic electron microscopy (cryo-EM), intact protein mass spectrometry, and small-molecule high-resolution and tandem mass spectrometry to probe the catalytic mechanism of human CerS6.

## Results

### Cryo-EM structure of human CerS6

We recombinantly expressed the human CerS6 protein, truncated at Asp350 to remove the predicted disordered C-terminus, in suspension-adapted HEK293 cells using the BacMam system. The protein was purified in glyco-diosgenin (GDN), yielding a mixture of monomeric and dimeric species on size-exclusion chromatography (Extended Data Fig. 1a-d). While we selected the dimer species for structural studies to maximize particle size and possible symmetry for single-particle cryo-EM, its relatively low molecular weight (84 kDa) is still challenging for current methods. We therefore raised camelid nanobodies against the purified protein and solved the structure of a CerS6 dimer with one nanobody bound per monomer at a nominal resolution of 3.2 Å resolution (Fig. 1a,b; Extended Data Table 1; Extended Data Fig. 2). The CerS6-Nb22 complex was well resolved throughout the membrane-spanning region, but the Hox-like domain was less well-defined. This allowed for unambiguous tracing of CerS6 and Nb22 with the exception of the Hox-like domains, where residues 72-119 were fitted into the envelope of the cryo-EM map as rigid bodies (see Methods) (Extended Data Fig. 2e; Extended Data Fig. 3a).

**Figure 1.**
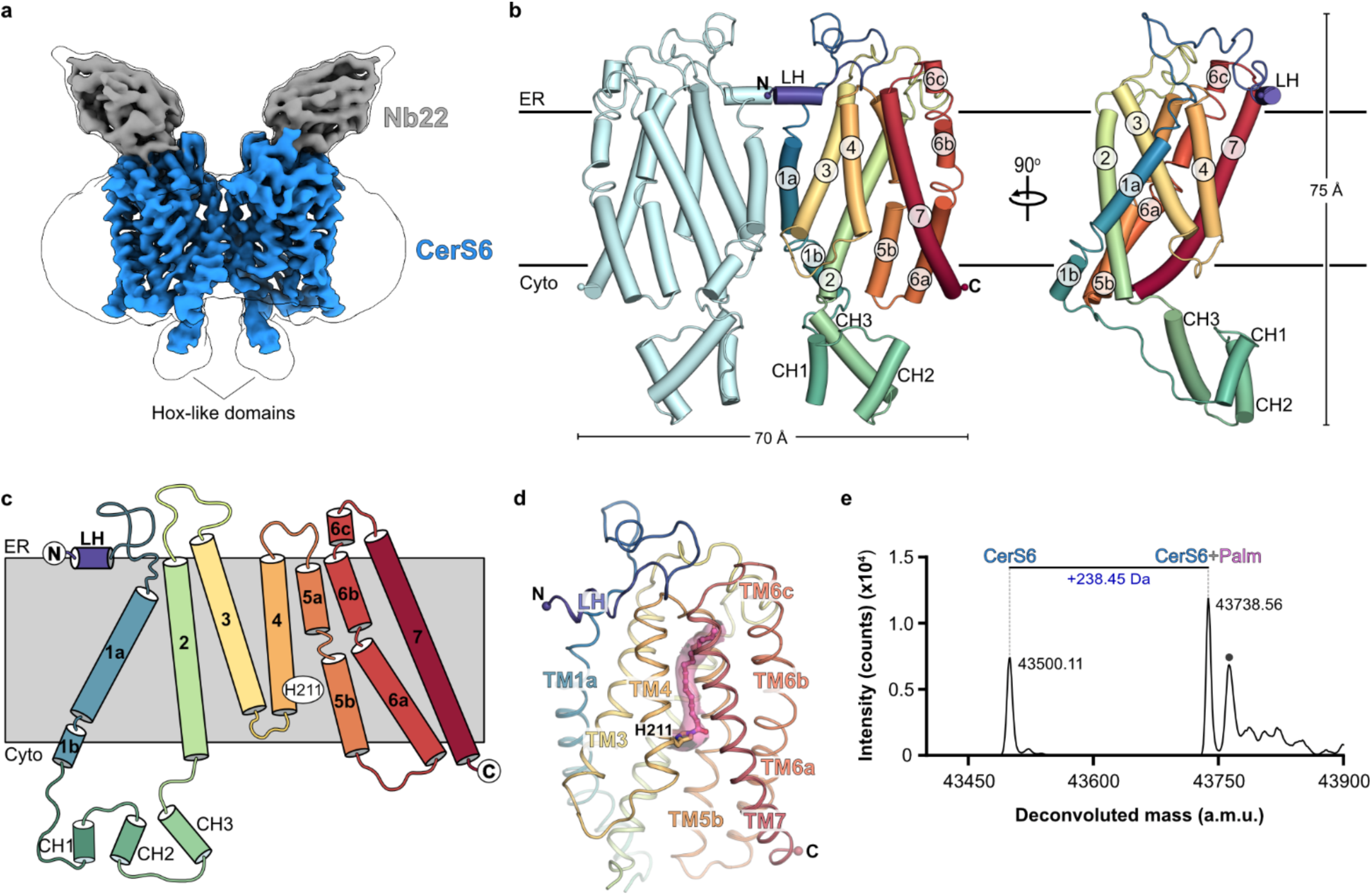
Cryo-EM structure of human CerS6. **a**, Cryo-EM map of the CerS6 dimer (blue) with one copy of Nb22 (gray) bound to each CerS6 monomer. **b,** Overall cartoon cylinder representation of the CerS6 dimer structure. One of the monomers is rainbow-coloured from purple (N-terminus) to red (C-terminus). **c,** schematic representation of the CerS6 7-TM helix topology. **d,** Cartoon representation of the transmembrane domain of a CerS6 monomer. The covalent acyl-imidazole species is shown in stick representation (acyl chain: pink carbon atoms), and the Coulombic potential map for this covalent species is shown as a pink transparent surface. The Hox-like domain was omitted for clarity. **e,** Denaturing intact protein MS analysis of purified CerS6 protein, revealing the presence of a covalent modification (+238.45 Da) which matches the expected mass shift corresponding to covalent attachment of a palmitoyl (Palm; C16:0) chain. This adduct peak was present in all purifications tested (n=6).

The cryo-EM structure reveals a CerS6 homodimer with one nanobody bound per monomer on the luminal face. Each monomer has 7 transmembrane helices (Fig. 1b,c), with the dimer interface formed by TM1a, the C-terminal end of TM3, and the TM3-TM4 loop. The TLC domain (TM2-7) forms a barrel containing two 3-TM units arranged as inverted repeats (Extended Data Fig. 4a), while TM1 is attached to the outside of the barrel through contacts with TM2/3. The TM2-7 barrel is assembled around a central cavity, which is open to the cytoplasm and closed to the ER lumen by a loop between TM6-7. This central cavity has a large cytoplasmic opening (approximately 23 x 12 Å), creating a vestibule that funnels towards the midpoint of the membrane, where it forms a ∼5 Å wide tunnel that stretches towards the ER side.

Interestingly, the fold of the TM2-7 barrel of CerS6 resembles that of the 6-TM barrels found in very-long-chain fatty acid elongase ELOVL7^34^ and TMEM120A/B^35,36^, as identified by DALI^37^. Additionally, a structurally homologous 6-TM barrel is predicted in the AlphaFold2 models of 3-hydroxyacyl-CoA dehydratases (HACD1-4) (Extended Data Fig. 4b), which participate in the same four-enzyme acyl-CoA elongation cycle as ELOVLs^38^. The observation that different acyl-CoA-modifying enzyme families share a similar 6-TM barrel structure is intriguing, suggesting that this architecture is likely optimized for the specific recognition and reaction with acyl-CoA substrates.

### CerS6 co-purifies with a covalently bound C16:0 acyl chain

Surprisingly, the cryo-EM density map revealed the presence of a long density extending from near the membrane mid-point to the occluded ER end of the central cavity, spanning the entire length of the narrow tunnel (Fig. 1d; Fig. 2a). This density was continuous between protein and the unknown molecule, suggesting a covalent modification by a lipid. To probe the presence of covalent adducts, we performed denaturing intact protein mass spectrometry (MS) analysis of purified protein samples, and identified a +238.45 Da mass shift (Fig. 1e and Extended Data Fig. 1f,g). Notably, this adduct mass agrees with a covalently attached C16:0 acyl chain (theoretical: +238.41 Da). Strikingly, the shape and length of the density in the cryo-EM map is also consistent with a bound C16 acyl chain. A covalent bond is seen linking the acyl chain to the imidazole Nε of His211 (TM4) (Fig. 2b; Extended Data Fig. 3b), an absolutely conserved histidine that is required for CerS activity^17,18^. Thus, based on the extended density in the cryo-EM map and the observed mass adduct, we have modelled a palmitoyl (C16:0) chain covalently attached to His211.

**Figure 2.**
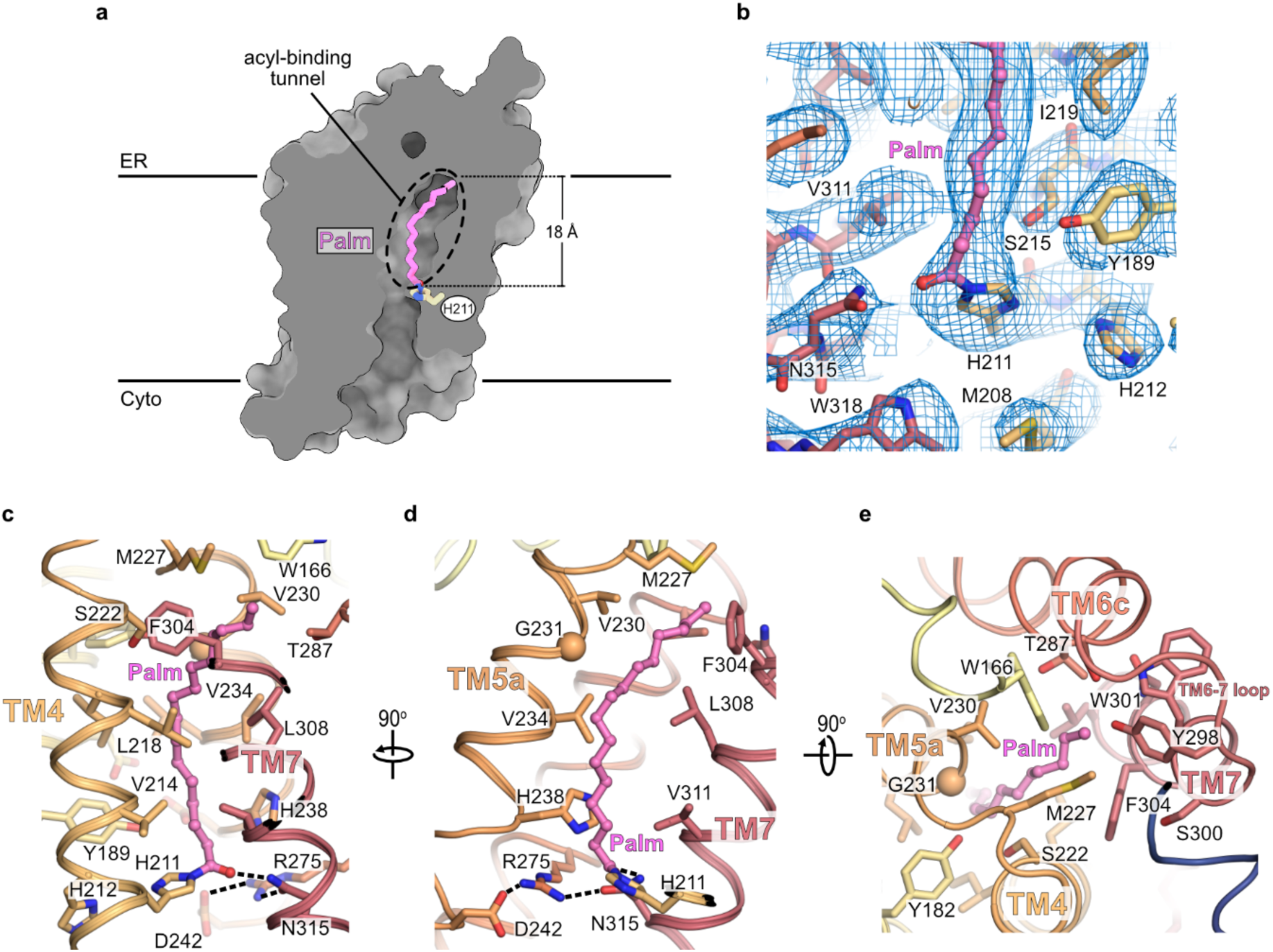
CerS6 contains an acyl chain-binding tunnel buried deep in the membrane. **a**, Cutaway molecular surface representation of the CerS6 transmembrane region, revealing the presence of a long central cavity spanning the entire length of the protein. The palmitoyl chain covalently attached to His211 is bound in a narrow tunnel on the ER luminal half of the central cavity. **b**, Coulombic potential map in the region around the covalent linkage of the acyl chain to His211. **c-e**, CerS6’s acyl-binding tunnel, viewed from the **(c,d)** membrane plane or **(e)** from the ER face. Side-chains lining and capping the tunnel are shown as sticks. Hydrogen bonds in the active site are shown as dashed lines.

The acyl chain is surrounded in the narrow tunnel primarily by hydrophobic side chains from TM4, TM5 and TM7 (Fig. 2c-e). The tunnel is sealed at the ER end by the TM6-7 loop (Fig. 2e), revealing how the chain-length preference of CerS6 for C16-CoA is determined by limiting the number of carbons that can fit in the acyl-binding tunnel. This observation is in agreement with earlier reports that this loop determines CerS acyl chain specificity^22^.

### Catalysis by CerS6 proceeds via a ping-pong mechanism

Acyltransferases catalyze two-substrate, two-product reactions that can proceed via either ternary-complex or double-displacement (ping-pong) mechanisms^39^. In a ternary-complex mechanism, both the acyl donor and acceptor bind simultaneously and the enzyme catalyzes the direct acyl transfer from one substrate to the other. In contrast, a ping-pong mechanism involves two independent steps. Initially, reaction with the first substrate (the acyl donor) results in the transfer of the acyl chain to a nucleophilic residue, forming a covalent acyl-enzyme intermediate. This acyl chain is then transferred to the second substrate (the acyl acceptor) in the second step of the reaction.

The CerS6 structure reveals that there is not enough space for both substrates to bind in the central cavity and access the active site at the same time, thereby indicating a ternary-complex mechanism is unlikely. Furthermore, the C16:0 chain covalently attached to the conserved His211 fits with CerS6’s selectivity for palmitoyl-CoA and suggests that this species could correspond to the acyl-enzyme intermediate of a ping-pong type reaction mechanism. To investigate whether the acylated enzyme species corresponds to a real catalytic intermediate, we incubated the purified protein with the enzyme’s second substrate, the long-chain base sphinganine (Fig. 3a). Using denaturing intact protein MS, we observed the complete loss of the C16:0-modified protein after sphinganine addition (Fig. 3b and Extended Data Fig. 5a). Furthermore, high-resolution mass spectrometry (HRMS) revealed that incubation of the acylated protein with sphinganine led to the production of C16:0 dihydroceramide (observed *m/z*: 540.5380; theoretical *m/z*: 540.5350; mass error: +5.55 ppm) (Fig. 3c). Overall, these results clearly demonstrate that sphinganine reacts with the acyl-enzyme intermediate to de-acylate the enzyme and generate the expected reaction product of dihydroceramide, thus supporting a ping-pong reaction mechanism for CerS6. Taken together with the structural data, this implies that His211 acts as the nucleophile in the first step of the reaction.

**Figure 3.**
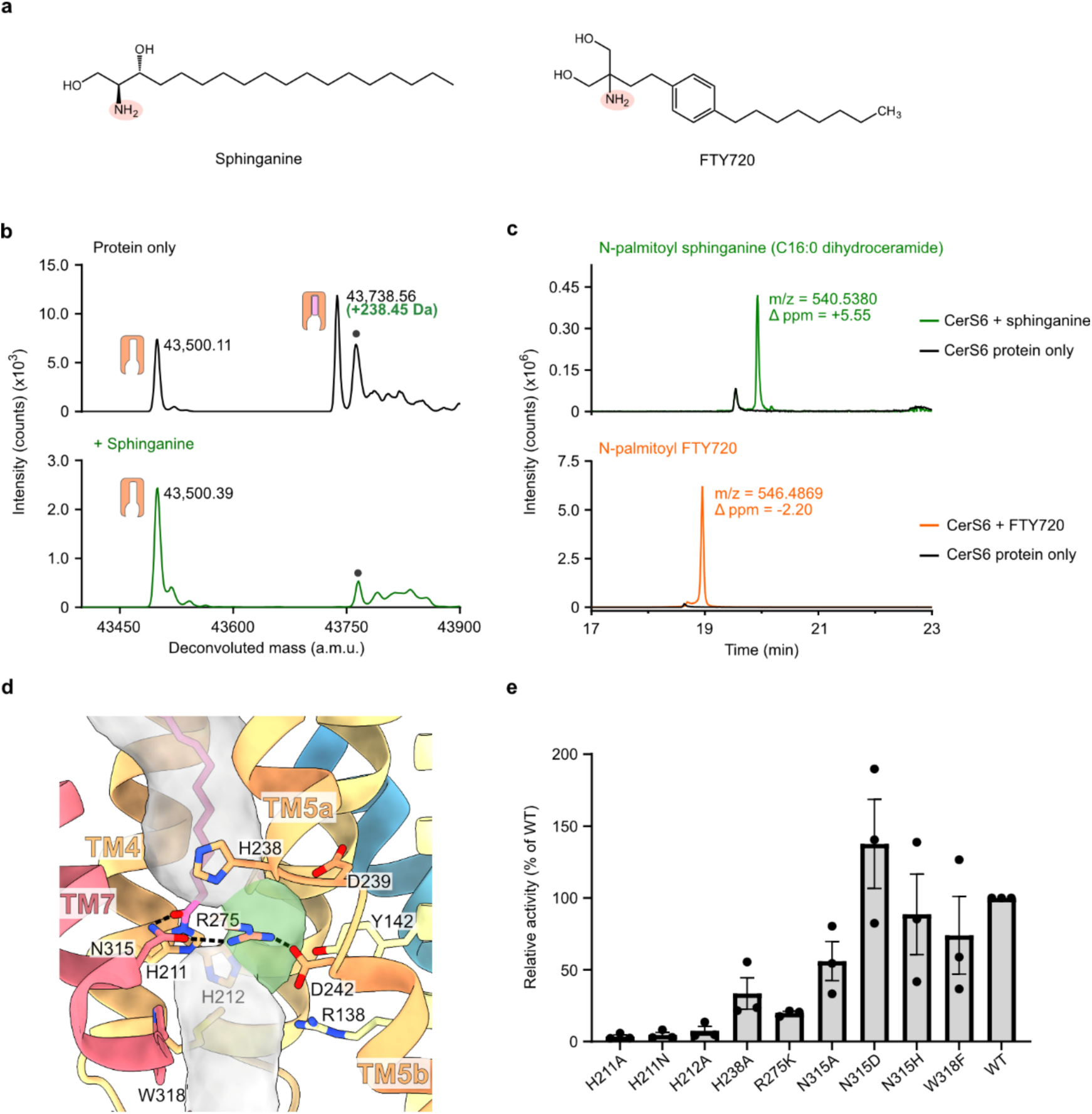
Unraveling the catalytic mechanism of CerS6 using mass spectrometry. **a**, chemical structures of the natural long-chain base substrate sphinganine (dihydrosphingosine) and the drug FTY720 (fingolimod, Gilenya). The homologous primary amines are highlighted in salmon. **b**, Intact mass analysis of protein samples after incubation in the absence of substrates (black) or in the presence of sphinganine (green) (n = 3 biological replicates; see Extended Data Fig. 5 for replicate traces). Deconvoluted mass peaks are indicated as follows: unmodified enzyme, orange icon; covalent acyl-enzyme species, orange and pink icon; background species present in all traces, gray circle. **c**, LC-HRMS detection of the reaction products. Extracted ion chromatograms (EICs) for the expected [M+H]^+^ ions of each of the reaction products after incubation of the acyl-enzyme intermediate with sphinganine (C16:0 dihydroceramide, green) or FTY720 (N-palmitoyl FTY720, orange). EICs obtained after incubation in the absence of substrates are shown in black. **d**, CerS6’s active site, viewed from the plane of the membrane. The central cavity is shown as a transparent gray surface, highlighting the presence of a side pocket (highlighted in green) adjacent to the acyl carbonyl.

Hydrogen bonds are shown as dashed lines. **e**, Mutational analysis of the CerS6 active site by comparison of the C16:0 dihydroceramide synthase activity of wild-type (WT) and active site mutants. n=3 independent biological replicates. Gray bars correspond to the mean of the biological replicates. Data points represent each biological replicate, corresponding to the mean of four technical replicates. Error bars show the SEM.

Notably, the CerS substrate sphinganine and FTY720 are chemically similar (Fig. 3a), with potential clinical consequences to drug pharmacokinetics as 15% of FTY720 becomes N-acylated in human subjects^40^. Although the major N-acyl FTY720 metabolites identified were N-stearoyl and N-2-hydroxystearoyl FTY720, there is evidence suggesting that a small amount of N-palmitoyl (C16:0) FTY720 is formed^40^. Which enzyme catalyzes the N-acylation of FTY720 has not been determined, but a putative role for CerS enzymes or the reverse reaction of ceramidases has been proposed^40^. Incubation of acyl-CerS6 with FTY720 results in a small increase in the protein’s thermostability, suggesting it binds to the purified enzyme (Extended Data Fig. 1e). To investigate whether purified CerS6 can N-acylate FTY720 *in vitro*, we incubated the enzyme with this compound, and subsequently identified C16:0 FTY720 in the reaction mixture (observed *m/z*: 546.4869; theoretical *m/z*: 546.4881; mass error: -2.20 ppm) (Fig. 3c), the expected product of a reaction between FTY720 and the CerS6 bound acyl chain. The proposed structure of this C16:0 FTY720 reaction product was further validated based on the product ion MS/MS spectrum of its [M+H]^+^ ion (Extended Data Fig. 6). However, no obvious loss of the acyl-enzyme mass peak was observed when the reaction mixture was analyzed by denaturing intact protein MS (Extended Data Fig. 5b), suggesting that FTY720 is a poor acyl acceptor for acylated CerS6.

### Highly conserved residues line the catalytic site

CerS6’s Lag1p motif (Arg202-Tyr253) (TM4-5) contains the highly conserved His211/His212 (TM4) and Asp239/Asp242 (L5a-b/TM5), which have been previously suggested to be located in the catalytic site and have been shown to be essential for function in mammalian CerS^17^ and *S. cerevisiae* Lag1p^18^.

Examining the CerS6 structure, we noted these residues line the central cavity near the mid-point of the membrane (Fig. 3d). Notably, the co-purified covalently linked C16:0 chain is covalently attached to the first histidine of the Lag1p motif (His211) (Fig. 2a-d), which appears to act as the nucleophile in the first step of the reaction and is required for CerS6 function (Fig. 3e). Although the observation that the mechanism of CerS6 involves a nucleophilic histidine was surprising - as cysteines or serines are more commonly employed as nucleophiles - there are multiple well-documented examples of other enzymes using nucleophilic histidines in their catalytic mechanisms^34,41–45^.

Histidine nucleophiles are typically oriented and activated by hydrogen-bonding between a proton at Nδ of the imidazole ring and a hydrogen bond acceptor, ensuring that Nε remains unprotonated^34,42,46^. Examining our structure, we noted the proximity of His212 to the acylated His211 in CerS6, which immediately suggested this residue is the necessary hydrogen bond acceptor. While the Nδ of His212 is located 3.84 Å away from the Nδ of His211, it is plausible that the two side chains may interact in the acyl-CoA-bound state. Indeed, we found that mutation of His212 to alanine results in the complete loss of activity (Fig. 3e). Residues equivalent to His212 of CerS6 are also essential in the homologous mouse and human CerS1^17,47^ and yeast Lag1p^18^. This demonstrates the second histidine plays a key role in catalysis, and explains the pathogenic effect of mutating the equivalent His of CerS1 to a Gln in patients with progressive myoclonic epilepsy and dementia^47^. Moreover, we have recently demonstrated that the reaction of human ELOVL7 with acyl-CoA substrates also involves a double histidine motif, where the first histidine acts as a nucleophile and is activated by hydrogen-bonding to the second^34^. Strikingly, structural alignment of CerS6 and ELOVL7 places the histidine pairs in equivalent positions (Extended Data Fig. 4c), suggesting that these residues may play similar roles in the first step of their respective reactions.

Formation of the acyl-enzyme intermediate proceeds via an oxyanion transition state, which is usually stabilized by hydrogen bonding. Within the acyl-CerS6 structure, the carbonyl oxygen of the covalently attached C16:0 chain is within hydrogen-bonding distance (2.8 Å) of the Nδ of Asn315 (TM7). Based on the proximity of these moieties, we hypothesized that this interaction could stabilize the formation of the oxyanion transition state during catalysis. Consistent with this notion, an Asn is present at the equivalent position in CerS1, and CerS2-5, Lag1p and Lac1p have a His at this position (Extended Data Fig. 7) which may likewise act as a hydrogen-bond donor. Indeed, we found that an Asn315His mutant preserves activity. Surprisingly, however, mutation of Asn315 to either an Ala or an Asp did not result in loss of activity (Fig. 3e). It is unclear if the hydrogen bond in the wild-type structure is not required for function, or if these mutations allow a water molecule to enter the reaction site and act as a substitute hydrogen bond acceptor. Asn315 is involved in a hydrogen-bonding network involving the conserved Arg275 (L6a-b) and Asp242 (TM5b). We found that an Arg275Lys mutant retained only approximately 20 % of wild-type activity (Fig. 3e), suggesting that the interactions established by the guanidino group of Arg275 to align neighbouring residues in the active site are important for function.

There is a third histidine on TM5a which is conserved in mammalian CerS but is a methionine in yeast Lag1p and Lac1p, and is therefore of ambiguous importance to the enzymes’ reaction cycle. In the CerS6 structure this residue, His238, is located above the acyl-imidazole carbonyl and lays against the hydrocarbon chain (Fig. 3d). We found that mutation of this His238 to alanine reduced activity by over 50 % (Fig. 3e), and therefore suggest this residue may participate in aligning the acyl chain.

Finally, we noticed the presence of a side pocket in the active site of CerS6, adjacent to His211 (Fig. 3d) and lined by Tyr142 (TM2), Tyr189 (TM3), His212 (TM4), Asp239 (L5a-b), Asp242 (TM5b) and Arg275 (L6a-b). The position and chemistry of this pocket relative to the carbonyl of the acyl-enzyme intermediate suggests this is likely to be the site where the amino alcohol moiety of the second (long-chain base) substrate binds. Supporting this notion, the residues of this pocket are highly conserved (Extended Data Fig. 7). This pocket therefore likely ensures the primary amine of the long-chain base is ideally located for a nucleophilic attack on the carbonyl carbon of the acyl-imidazole intermediate in the second step of the reaction.

### Structure of N-acyl fumonisin B_1_-bound CerS6

Having captured the intermediate state of the CerS6 enzyme, we next set out to examine the structural basis of CerS inhibition by FB_1_. To this end, we incubated purified acyl-CerS6 with fumonisin B_1_, and determined the complex’s structure to a nominal resolution of 2.95 Å (Fig. 4a,b and Extended Data Fig. 8). In comparison with the covalent intermediate state, the overall structure of each CerS6 monomer did not change (transmembrane region backbone RMSD = 0.39 Å), although the flexible Hox-like domains rotated by approximately 10°.

**Figure 4.**
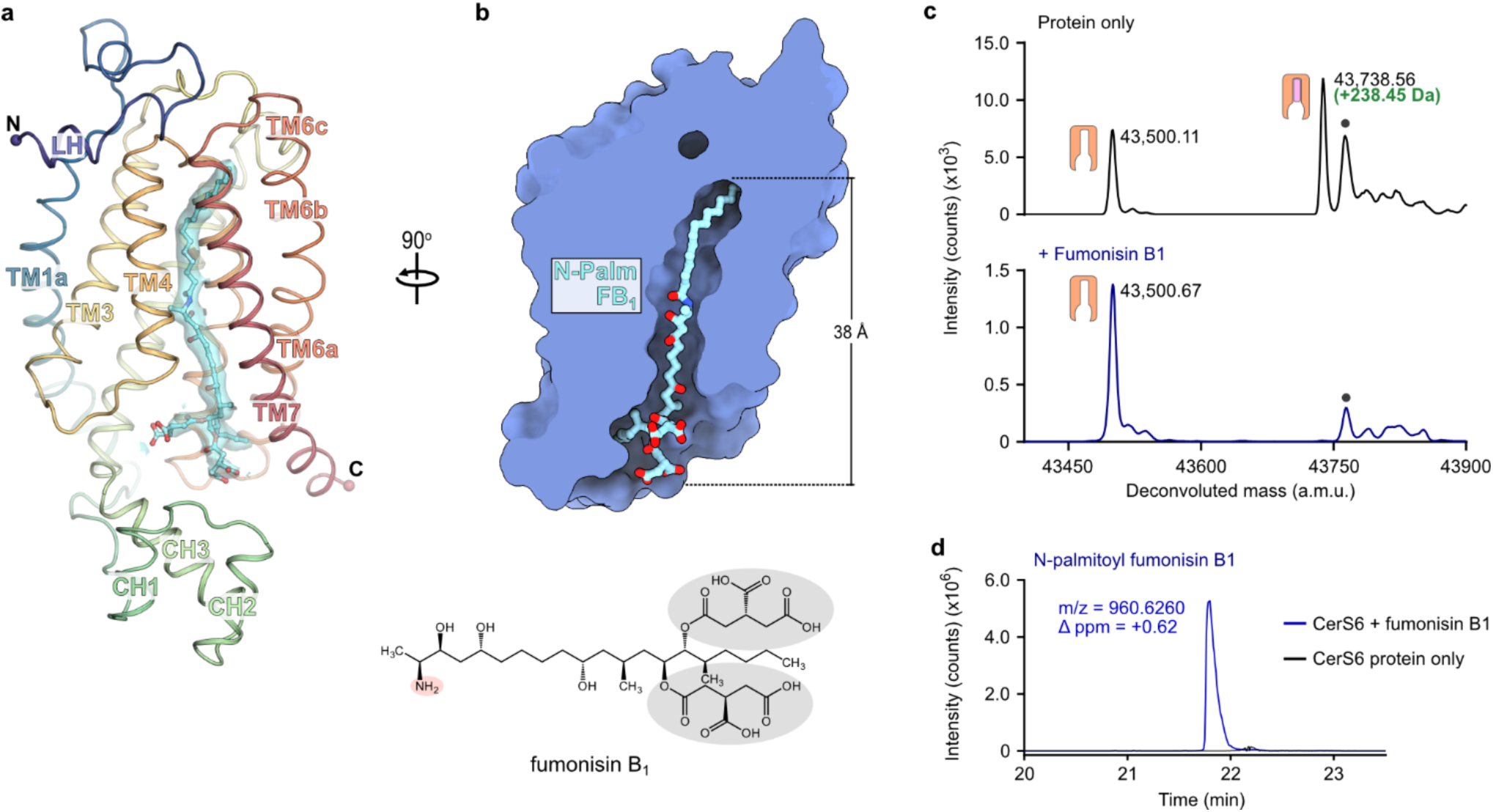
Cryo-EM structure of CerS6 in complex with N-palmitoyl fumonisin B_1_. **a**, Cartoon representation of CerS6 with bound N-palmitoyl fumonisin B_1_ (shown as sticks; cyan carbon atoms). The cryo-EM density of the bound product is shown as a transparent cyan surface. **b**, Cutaway molecular surface representation, revealing that the N-palmitoyl fumonisin B_1_ species occupies the entire length of the central cavity. The Hox-like domain has been omitted for clarity. **c**, Intact mass analysis of protein samples after incubation in the absence of substrates (black) or in the presence of the mycotoxin fumonisin B_1_ (blue) (n = 3 biological replicates; see Extended Data Fig. 5 for replicate traces). **d,** LC-HRMS detection of N-palmitoyl fumonisin B_1_. The EIC for its expected [M+H]^+^ ion is shown after incubation of the acyl-enzyme intermediate with fumonisin B_1_ (blue) or in the absence of the toxin (black). *Inset:* chemical structure of fumonisin B_1_. Its primary amine (salmon circle) and tricarballylic acid groups (gray circles) are highlighted.

The cryo-EM map of the putative FB_1_-bound complex contains a long, continuous non-protein density starting within the ER luminal end of the narrow tunnel and extending to the cytoplasmic entrance to the central cavity (Fig. 4a and Extended Data Fig. 3c,d). The observed density is too long to be FB_1_ alone. Notably, incubation of palmitoyl-CerS6 with FB_1_ completely removed the acyl group from the enzyme (Fig. 4c and Extended Data Fig. 5a and generated C16:0-fumonisin B_1_ (observed *m/z*: 960.6260; theoretical *m/z*: 960.6254; mass error: +0.62 ppm) (Fig. 4d). Noting the covalent bond between the enzyme and the palmitoyl group has been replaced by a bond to the toxin, we modeled this density as C16:0-FB_1_ (Fig. 5a and Extended Data Fig. 3c). These results indicate FB_1_ has reacted with the covalent acyl-enzyme intermediate, releasing His211 and generating the N-acyl FB_1_ product. This product remains bound to the enzyme, thus acting as an inhibitor. This finding is consistent with earlier reports that CerS enzymes can N-acylate FB_1_ in cultured cells and in rats^48,49^.

**Figure 5.**
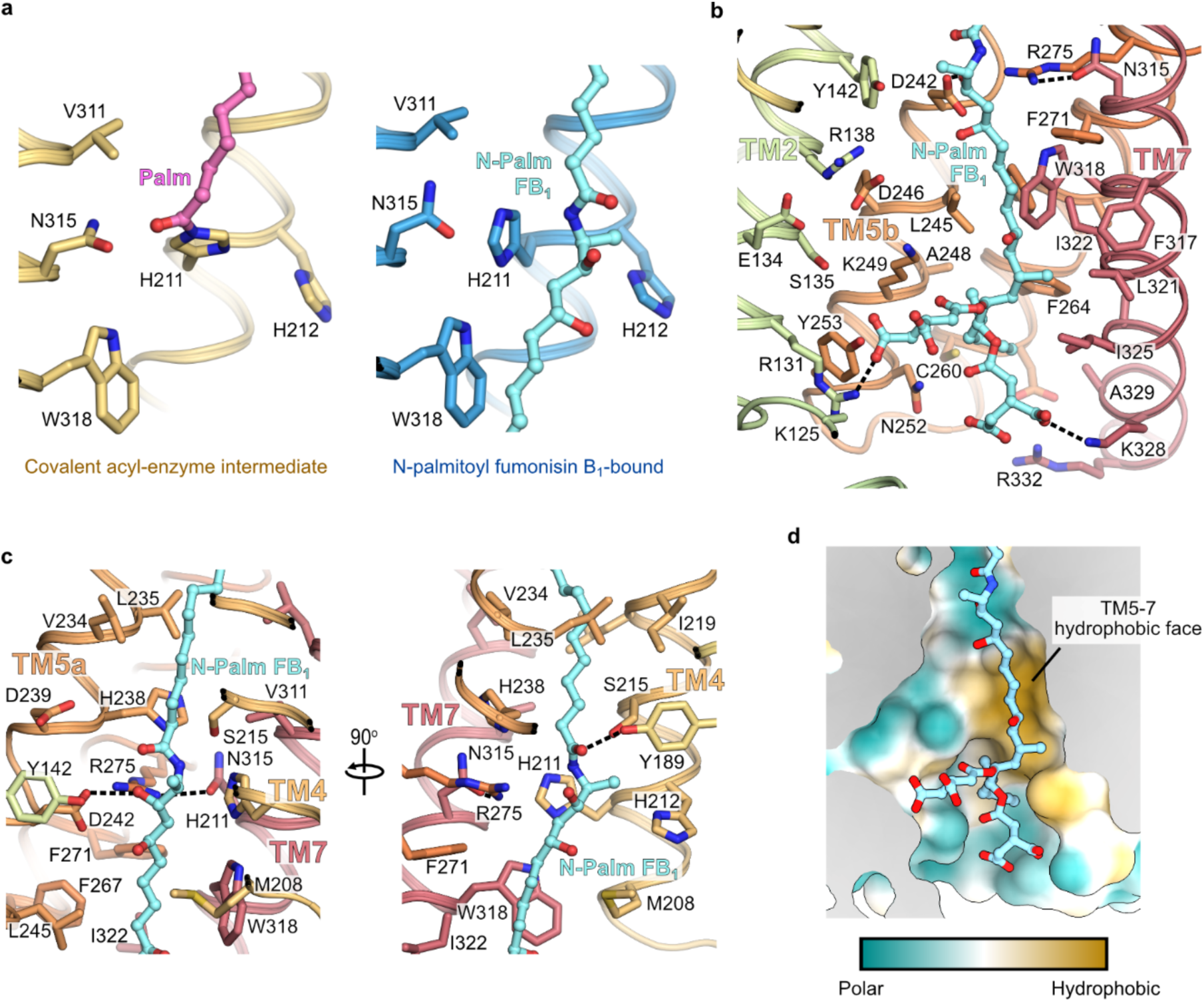
Binding mode of N-palmitoyl fumonisin B_1_. **a**, Close-up view of the CerS6 active site in the covalent acyl-enzyme intermediate and N-palmitoyl FB_1_-bound states, showing the transfer of the palmitoyl chain from His211 to the toxin. **b**, Cytoplasmic portion of the central cavity, viewed from the membrane plane. Residues lining the cavity are shown as sticks. Polar interactions between the carboxylates of the tricarballylic acid groups of fumonisin B_1_ and positively charged residues on TM2 and TM7 are shown as dashed lines. **c**, Active site, viewed from the plane of the membrane. **d**, Polar and non-polar surfaces on the cytoplasmic half of the central cavity. Cutaway molecular surface view, showing that the hydrocarbon chain of fumonisin B_1_ interacts with the large hydrophobic face formed by TM5-7.

The N-acyl FB_1_ species spans the entire length of the central cavity, with the acyl chain remaining bound in the narrow tunnel at the occluded ER end of the cavity, and the FB_1_ portion sitting in the wider cytoplasmic entrance (Fig. 4b and Fig. 5b-c). The newly formed amide bond sits in the active site (Fig. 5c), with the carbonyl oxygen forming a hydrogen bond to Tyr189 (TM3), and the amide NH hydrogen bonding to the Nε of His211 (TM4). The 3-hydroxyl group of FB_1_, which is also present in sphingoid base substrates, forms a hydrogen bond with the strictly conserved and catalytically essential Asp242 (TM5b)^17,18^. Such an interaction with the 3-hydroxyl group of sphingoid base substrates would optimally place the sphingoid primary amine to attack the carbonyl carbon. However, further work is required to investigate this hypothesis.

Below the active site, the cytoplasmic vestibule of the central cavity has both polar and non-polar surfaces, where we noted that the hydrocarbon chain of FB_1_ interacts with the long hydrophobic face formed by TM5-7 (Fig. 5b,d). This hydrocarbon chain lies against the conserved Trp318 of TM7 just beneath the active site. Mutation of the corresponding residue in CerS5 to an alanine has been previously shown to abolish enzymatic activity^50^. In contrast, introducing a phenylalanine at this position by a W318F mutation in CerS6 retained approximately wild-type levels of activity (Fig. 3e). From these results, we hypothesize the conserved Trp orients the sphingoid base substrate to place its amino alcohol moiety in the side pocket within the active site for the second step of the reaction.

The face of the central cavity opposite the FB_1_ hydrocarbon binding surface displays a polar character, and is lined by conserved charged residues that may participate in CoA binding (Fig. 5b,d). Accordingly, the positive charges provided by Lys125 and Arg131, located at the cytoplasmic entrance to the cavity, adjacent to the Hox-like domain, have been previously demonstrated to be important for the function of CerS5 and CerS6^20^. Furthermore, mutation of Lys131 in CerS3, equivalent to CerS6’s Arg131, to alanine has been implicated in autosomal recessive congenital ichthyosis (ARCI)^51^, a disorder of epidermal keratinization which can be caused by loss of ≥ C26 ceramides due to mutations in *CERS3*^52,53^.

The two tricarballylic acid (TCA) groups of FB_1_ span the width of the cytoplasmic end of the central cavity, where one of the groups interacts with Arg131 (TM2) and the other with Lys328 (TM7) (Fig. 5b). Thus, through these charge-charge interactions, FB_1_ links the first helix of the first 3-TM bundle and the last helix of the second. Supporting the notion that FB_1_ acts to restrain the structure, atomistic molecular dynamics (MD) simulations of covalent intermediate and N-acyl FB_1_-bound CerS6 monomers indicate that a bound N-palmitoyl FB_1_ reduces the overall flexibility of the protein, including TM6, TM7, and the Hox-like domain (Extended Data Fig. 9a-c). Moreover, while Lys125 is not resolved in the cryo-EM maps, in our simulations Lys125 forms a salt bridge with the same TCA group that contacts Arg131 (Extended Data Fig. 9d). Further, the simulations show that this TCA also contacts Lys249 (TM5b), Tyr253 (TM5b), as well as Arg118 and Arg121 of the last helix of the Hox-like domain. Therefore, FB_1_ appears to mimic a number of CoA interactions through the binding of one of its TCA moieties at the polar face of the cytoplasmic cavity. Finally, on the opposite side of the cavity, the other TCA moiety can form salt bridges with Lys328 or Arg332 (TM7) (Extended Data Fig. 9d). Overall, our cryo-EM structure and simulations reveal that FB_1_ is anchored by polar interactions at opposite sides of the 6TM barrel, which likely hinder product release, arresting the enzyme in an inhibitory product-bound state.

### Bound lipid

In examining our CerS6 cryo-EM maps, we noted both conditions clearly contain an ordered non-protein density near the dimer interface, adjacent to the LH-TM1 loop and TM4 and likely corresponding to a bound lipid. There, we modelled a phosphatidylcholine molecule, as its head group is consistent with the shape, size and environment of the observed density (Extended Data Fig. 3e). The choline head group is nestled between conserved Trp9 and Trp21 in a classic aromatic box for binding quaternary amines^54^. Adjacent to the choline-binding site is Trp14, which is strictly conserved in CerS2-6. Interestingly, mutation of the corresponding tryptophan residue in CerS3 to an arginine has been found in patients with ARCI, and the mutant protein was found to be catalytically inactive^52^. In the CerS6 structure, Trp14 does not directly contact the choline head group, but it could play a structural role in forming the lipid-binding site.

## Discussion

Ceramide synthases carry out an essential step in sphingolipid biosynthesis and are promising drug targets. Yet, our limited understanding of their structures and biochemical mechanism has hindered their therapeutic targeting. Here, we provide structural snapshots of CerS6 at two stages of its reaction cycle, revealing its ping-pong (double-displacement) reaction mechanism which uses a histidine nucleophile that attacks the acyl-CoA thioester to form a stable acyl-imidazole intermediate. This intermediate subsequently reacts with the sphingoid long-chain base substrate to yield the final ceramide or dihydroceramide reaction product (Fig. 6).

**Figure 6.**
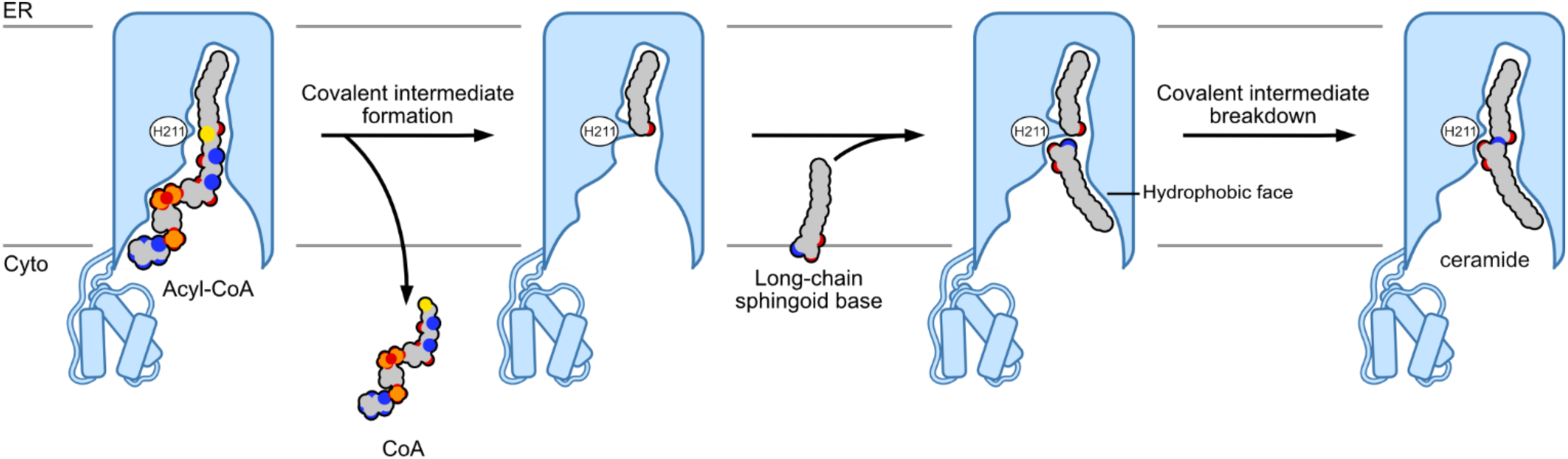
Proposed double-displacement (ping-pong) mechanism of ceramide synthases. Initially, the acyl-CoA substrate binds with the acyl chain buried deep within the central tunnel and the CoA moiety sitting near the cytoplasmic entrance to the central cavity. In the first step, the nucleophilic attack of His211 on the acyl-CoA thioester carbonyl results in thioester cleavage, covalent acyl-imidazole intermediate formation, and release of CoA. Subsequently, the long-chain sphingoid base substrate binds, with its hydrocarbon chain interacting with the hydrophobic face of the central cavity and its amino alcohol moiety sitting in the side pocket in the active site. In the second step of the reaction, the primary amine of the long-chain base attacks the acyl-imidazole intermediate, leading to covalent intermediate breakdown and formation of the final N-acyl sphingoid base (ceramide) product.

Although CoA and the hydrocarbon chain of long-chain bases likely interact at distinct faces of the cytoplasmic vestibule, there is not enough space for the acyl-CoA and long-chain base substrates to bind at the same time. Therefore, upon thioester cleavage and covalent intermediate formation, the CoA product needs to exit the central cavity before the long-chain base substrate can bind. We hypothesize that the amino alcohol group of the long-chain base would bind in a discrete side pocket lined by polar residues conserved across the entire CerS family. This long-chain base is further oriented for the second step of the reaction by hydrophobic residues located at the entrance to this pocket, particularly by the conserved Trp318 (TM7). The primary amine of the long-chain base then attacks the acyl-enzyme intermediate to form the final reaction product. Release of this product into the membrane likely occurs between two transmembrane helices, and the two halves of the 6TM barrel immediately suggest an egress route between TM4 and TM7. Indeed, we found that the TCA groups of N-palmitoyl FB_1_ attach to both halves of the barrel and restrict the movement of TM7, suggesting these interactions likely prevent that product’s release. Indeed, this proposed role for the TCA groups of FB_1_ is consistent with previous reports that the hydrolysed form of FB_1_, lacking the TCA moieties, is a weaker CerS inhibitor^55^. However, FB_1_ is competitive towards both substrates^30^. This suggests FB_1_’s mode of inhibition of CerS is likely to be multifaceted, sterically blocking substrate binding to the apo state, and locking the protein in a product bound state as shown here.

Recently, Zelnik et al. postulated that CerS2, and ceramide synthases more generally, use a ternary complex reaction mechanism^50^. In this model, the substrates bind orthogonally, and the long-chain base would access the active site through a side entrance located near the membrane mid-point, where it would bind parallel to the membrane plane. This proposed substrate-binding arrangement is similar to that of small-molecule membrane-bound O-acyltransferases (MBOATs) and led to the suggestion that CerS may catalyze the direct transfer of the acyl chain from acyl-CoA to the long-chain base in a single step involving the two active site histidines^50^. In addition, while this manuscript was under review, the cryo-EM structure of yeast Lac1p bound to an acyl-CoA was reported. That structure revealed a small lateral opening in Lac1p adjacent to the active site^56^. This observation led the authors to hypothesize that it could potentially correspond to the entry route for sphingoid base substrates^56^. However, this lateral opening is not present in our cryo-EM structures of CerS6 or in our molecular dynamics simulations. Instead, our experimental structures and biochemical studies of CerS6 present several pieces of evidence which support a ping-pong type mechanism, and challenge a ternary complex reaction. First, CerS possess a completely different fold to that of the MBOATs. Indeed, CerS6’s structure is strikingly similar to the fatty acid elongase ELOVL7, forming a 6-TM barrel that encloses a narrow tunnel with an essential and structurally superimposable active site histidine pair, which we have also recently shown to catalyze acyl chain transfer via a ping-pong reaction mechanism^34^. Further, purification of a stable acyl-CerS6 adduct species is incompatible with a ternary complex reaction mechanism. Rather, by incubating acyl-CerS6 with sphinganine, FB_1_ or FTY720, we show that this species is a *bona fide* reaction intermediate which results in the expected product upon exposure to the second substrate. Additionally, we captured the product-bound state in the N-acyl FB_1_-bound structure, where the toxin’s hydrocarbon chain binds perpendicularly to the membrane plane and interacts with the hydrophobic face of the central cavity formed by TM5-7. This binding mode in the N-acyl FB_1_-bound structure conflicts with a MBOAT-style substrate coordination, but is still consistent with the action of CerS on the fluorescently labelled NBD-sphinganine^57^ as the bulky NBD moiety would sit near the wide cytoplasmic entrance of the central cavity. Crucially, FB_1_ is a structural analog of sphinganine which undergoes N-acylation similarly to sphingoid base substrates, and its mode of inhibition is competitive towards sphinganine^30^, implying that FB_1_ and sphinganine likely bind to CerS in a similar manner. In aggregate, our data do not support a ternary complex reaction mechanism. Rather, an ELOVL-type ping-pong model is consistent with both our newly determined structures and biochemical results, and previously reported CerS mutants^17,18,47^.

Our observation that CerS6 can transfer a C16:0 acyl chain to the primary amine of FTY720 has important consequences for using compounds that mimic sphingoid bases in a clinical setting. The action of CerS partially explains the N-acylated FTY720 species found during the FTY720 drug trials^40^, and by extension this enzyme family may play a role in the pharmacokinetics of other sphingoid analogs. Further, our FB_1_ results reveal how this toxin can arrest the protein in a product-bound state (Extended Data Fig. 10), supporting the notion that unintended substrates of this enzyme family may result in the formation of inhibitory products. Taking advantage of this inhibition, ceramide or N-acyl FB_1_ mimetics could provide novel chemical scaffolds for new CerS inhibitors.

## Methods

### CerS6 cloning and expression

The *Homo sapiens CERS6* gene, encoding the CerS6 protein (Uniprot ID Q6ZMG9) was cloned into the baculovirus transfer vector pHTBV1.1-CT10H-SIII-LIC (adapted from the BacMam vector pHTBV1.1, provided by Frederick Boyce, Massachusetts General Hospital, Cambridge, MA) upstream of C-terminal TEV-cleavable 10x His and Twin-Strep tags. The construct used for structural determination lacked the final 42 amino acids, which are predicted to be disordered. For *in vitro* biotinylation, the avi-tagged CerS6 construct contained a GGGS linker between CerS6 and the Avi tag (GLNDIFEAQKIE), located upstream of the TEV cleavage site. Site-directed mutagenesis was performed using the Q5 site-directed mutagenesis kit (NEB).

Baculoviral DNA was generated by transposition of DH10Bac with the baculovirus transfer vector. Baculovirus was produced by transfecting *Sf*9 cells (Thermo Fisher Scientific) with the baculoviral DNA using the Insect GeneJuice transfection reagent (Merck Milipore). The virus was amplified by infecting *Sf*9 cells in the presence of 2 % fetal bovine serum and incubating for 72h at 27 °C. For large-scale protein expression in Expi293F GnTI-cells (Thermo Fisher Scientific) in Freestyle 293 expression medium (Thermo Fisher Scientific), cells were transduced with baculovirus and supplemented with 5 mM sodium butyrate. Cells were incubated in a humidity-controlled orbital shaker at 37 °C with 8 % CO_2_ for 48h, and subsequently harvested by centrifugation at 900 x g for 15 min. Pelleted cells were washed with phosphate buffered saline (PBS), pelleted again by centrifugation, flash-frozen in liquid N_2_ and stored at -80 °C. To compare mutant activity, wild-type and mutant proteins were expressed in Expi293F cells (Thermo Fisher Scientific) using the same protocol.

### Large-scale CerS6 purification

CerS6-overexpressing cells were re-suspended in Buffer A (50 mM HEPES pH 7.5, 200 mM NaCl, 5 % (v/v) glycerol) and solubilized with 1 % (w/v) lauryl maltose neopentyl glycol (LMNG) (Anatrace) and 0.1 % (w/v) cholesteryl hemisuccinate (CHS) (Sigma-Aldrich). Insoluble material was removed by centrifugation at 35,000 x g, and the protein was purified from the supernatant by batch binding to Strep-Tactin XT Superflow resin (IBA Lifesciences). The resin was washed initially with Buffer A containing 0.05 % (w/v) GDN, and subsequently with Buffer A supplemented with 1 mM ATP (Sigma-Aldrich), 10 mM MgCl_2_, and 0.05 % (w/v) GDN. CerS6 was eluted from the resin using Buffer A supplemented with 100 mM D-biotin (Fluorochem) and 0.02 % (w/v) GDN, and the C-terminal tag was cleaved by TEV protease digestion overnight. The His-tagged TEV protease was removed by binding to Co^2+^-charged TALON resin (Clontech) and the flow-through concentrated using a 100 kDa molecular weight cut off concentrator (Corning). The protein was finally purified by size-exclusion chromatography on a Superdex 200 Increase 10/300 GL column (GE Healthcare), pre-equilibrated in SEC buffer (20 mM HEPES pH 7.5, 200 mM NaCl, and 0.01 % (w/v) GDN).

For nanobody generation and selection, GDN was replaced in the purification buffers with 0.003 % (w/v) LMNG and 0.0003 % (w/v) CHS. For *in vitro* biotinylation of C-terminally avi-tagged CerS6, during the TEV cleavage step, the protein sample was further supplemented with 15 mM MgCl_2_, 15 mM ATP, 50 mM bicine pH 8.3, and BirA at a 1:15 (w/w) BirA:CerS ratio. Biotinylation efficiency was routinely monitored by denaturing intact protein mass spectrometry.

For structural determination, CerS6 was mixed with a 1.5 x molar excess of nanobody Nb22 and incubated for 1h on ice. The CerS6-Nb22 complex was purified by size-exclusion chromatography, concentrated to 5 mg mL^-1^ using a 100 kDa molecular weight cut off concentrator (Sartorius), and processed immediately for cryo-electron microscopy.

### Nanobody library generation and selection

Purified CerS6 was reconstituted at a 1:20 (protein:lipid (w/w) ratio into liposomes composed of POPC:POPG:DMPC:DMPG (7:3:7:3 (w/w)) as previously described^58^.

To obtain anti-CerS6 nanobodies, alpacas were immunized and the nanobody library was generated as previously described^59^, with the exception that 200 µg purified CerS6 in proteoliposomes were used for each immunization. All the procedures concerning alpaca immunization were approved by the Cantonal Veterinary Office of Zurich, Switzerland (License No. ZH 198/17). The resulting nanobody library was screened by biopanning against CerS6. Subsequently, 190 single clones from the enriched nanobody library were analyzed by ELISA for binding to CerS6. 96 ELISA-positive clones were Sanger sequenced and grouped in families according to their CDR length and sequence diversity^60^. Of these, 42 unique nanobodies were identified as belonging to 26 nanobody families and were taken for further validation.

Nanobodies were expressed in 50 mL scale in WK6 cells and purified from periplasmic extracts using Ni^2+^-NTA resin, as previously described^60^. Unique nanobodies were screened by biolayer interferometry (BLI) in an Octet Red 384 system (Sartorius) using Streptavidin SA biosensors (Sartorius) loaded with 40 µg mL^-1^ biotinylated CerS6 in SEC buffer containing 0.003 % (w/v) LMNG, 0.0003 % (w/v) CHS. Nanobodies with slow off rates were identified, and nanobody-22 was prioritized for structural studies.

### Thermal stability measurements

Thermal unfolding experiments were carried out on a Prometheus NT.48 instrument (NanoTemper Technologies) using 5 µM CerS6 solubilized in 0.003% LMNG and 0.0003% CHS. The protein was incubated for 1h at 4 °C in the presence or absence of 20 or 100 µM FTY720 (Sigma-Aldrich) or fumonisin B_1_ (Sigma-Aldrich). Proteins were heated from 20 °C to 95 °C, at a rate of 1 °C min^-1^, and unfolding was monitored by the ratio of fluorescence emission at 350 nm and 330 nm. The melting temperature was determined from the inflection point of the transition using the PR.ThermControl software (NanoTemper Technologies).

### Cryo-EM sample preparation and data acquisition

The purified CerS6-Nb22 complex (5 mg mL^-1^) was applied to freshly glow-discharged Quantifoil 200 mesh Au R1.2/1.3 grids. Plunge freezing in liquid ethane was carried out using a Vitrobot Mark IV (Thermo Fisher Scientific) set to 4 °C and 100 % humidity. For the CerS6-Nb22-FB_1_ complex, the SEC-purified CerS6-Nb22 complex was incubated with 120 µM fumonisin B_1_ (Sigma-Aldrich) for 10 minutes on ice prior to protein concentration and subsequently for an additional 2h on ice prior to grid preparation. EM grids were screened on a Glacios (OPIC, Oxford) and high resolution data were collected on a Titan Krios G3 microscope (LISCB, University of Leicester) operating at 300 kV and equipped with a Bioquantum Energy Filter (Gatan) (operated at 20 eV slit width) and a K3 direct electron detector (Gatan) at 130,000 x nominal magnification, in super-resolution mode (2x binning; physical pixel size: 0.656 Å px^-1^), using a defocus range between -0.8 µm and -2.4 µm. The total exposure dose was 56.3 e^-^ Å^-2^, fractionated over 50 frames. In total, 14,309 and 18,386 movies were collected for the CerS6-Nb22 and CerS6-Nb22-FB_1_ datasets, respectively.

### Cryo-EM data processing and model building

Movies were motion-corrected using MotionCor2^61^, and CTF estimation was carried out in cryoSPARC^62^ (version 3.3.1). Micrographs with bad CTF fitting (>5 Å) were excluded, yielding 14,196 and 16,647 micrographs for further analysis for the CerS6-Nb22 and for the CerS6-Nb22-FB_1_ datasets, respectively.

For the covalent acyl-enzyme intermediate dataset, particles were initially blob picked and extracted from a subset of micrographs, and representative well-resolved 2D classes were used for template-based picking of the entire dataset. 3,400,118 template-picked particles were extracted in a 360 px box Fourier cropped to 120 px corresponding to a pixel size of 1.968 Å px^-1^ and classified in two rounds of reference-free 2D classification yielding 1,261,928 good particles. Four a*b-initio* models were generated and used as reference models in heterogeneous refinement. Particles belonging to the well-resolved 2:2 CerS6-Nb dimer complex class (53 % of input particles) were selected and subjected to a further round of heterogeneous refinement. The best resolved set of particles (497,504) were then subjected to non-uniform refinement^63^ with C2 symmetry imposed, yielding a 3.21 Å reconstruction (FSC = 0.143). These aligned particles were imported into RELION using the csparc2star.py script from the UCSF pyem package^64^ and re-extracted in a 360 px box Fourier cropped to 270 px, corresponding to a pixel size of 0.875 Å px^-1^. In RELION, CTF parameters were refined, followed by Bayesian particle polishing^65^. The polished particles were subjected to 3D classification without image alignment (k =10, T=12). Certain classes displayed worse side-chain density and local deviations from C2 symmetry. Therefore, to improve map quality, the 93,680 particles belonging to the two highest-resolution C2-symmetric classes were re-imported into cryoSPARC, and subjected to a final non-uniform refinement with C2 symmetry, yielding a 3.22 Å reconstruction (FSC = 0.143). Importantly, even though CTF refinement, Bayesian polishing and 3D classification in RELION did not improve the nominal resolution, the final reconstruction reveals better defined features, including improved side-chain density in the CerS6 active site (Extended Data Fig. 2f,g).

The N-acyl FB_1_ dataset was processed using the same workflow as the covalent acyl-enzyme intermediate dataset, with the following changes. 3,998,935 template-picked particles were classified in two rounds of reference-free 2D classification, yielding 1,353,329 good particles. Four a*b-initio* models were generated and used as reference models in heterogeneous refinement. 641,462 particles belonging to the well-resolved 2:2 CerS6-Nb dimer complex class (48 % of input particles) were subjected to non-uniform refinement with C2 symmetry imposed, yielding a 3.17 Å reconstruction. After CTF refinement, Bayesian polishing and 3D classification without image alignment in RELION, the 154,239 particles belonging to the three highest-resolution C2-symmetric classes were selected. These particles were re-imported into cryoSPARC, and subjected to a final non-uniform refinement with C2 symmetry, yielding a 2.95 Å reconstruction (FSC = 0.143).

Atomic models were generated by fitting the CerS6 AlphaFold2^66^ prediction and a Phyre2^67^ nanobody homology model into the cryo-EM maps, and subsequently manually adjusted in Coot^68^. In both datasets, residues 72-119, corresponding to the Hox-like domains, were poorly resolved in the sharpened maps. However, their position is evident in unsharpened and blurred maps (Extended Data Fig. 2e; Extended Data Fig. 3a). In order to model this region, we used tight restraints to the Alphafold2 prediction for this domain and docked it into the envelope of the blurred maps (B_Blur_ = 200 Å^2^), and surface-exposed side-chains were truncated at Cβ. The atomic models were refined using PHENIX real-space refinement^69^ with secondary structure and Ramachandran restraints, and NCS constraints. Restraint dictionaries for FB_1_ and the covalently attached C16:0 chain were generated using AceDRG^70^. The final models comprise residues 2-330 (covalent intermediate state) or 2-334 (N-palmitoyl FB_1_-bound state) of CerS6, residues 1-124 of nanobody-22, the C16:0 chain covalently attached to His211 of CerS6 or the N-palmitoyl FB_1_ product, one phosphatidylcholine molecule and the first N-acetylglucosamine residue of the N-linked glycan visible on Asn18. Weighted Fo − Fc ligand difference maps were calculated using Servalcat ^71^, available in CCPEM^72^, by omitting the ligands from the atomic models.

### Denaturing intact protein mass spectrometry

The intact masses of purified protein samples were analyzed by denaturing intact protein mass spectrometry, conducted using an Agilent 1290 Infinity LC system in line with an Agilent 6530 Accurate-Mass quadrupole time-of-flight mass spectrometer (Agilent Technologies), as previously described^34^. Typically, 5-8 μg of purified protein (at 1.5-2.0 mg mL^-1^), diluted to 20 μL in 30 % methanol in 0.1 % formic acid, were used per injection. Data were acquired between 100-3200 *m/z* and analyzed using MassHunter Qualitative Analysis v.B.07.00 (Agilent Technologies) software. Peaks between 650-3200 *m/z* in the sum of the mass spectra obtained during protein elution were deconvoluted using the maximum entropy charge deconvolution algorithm.

The identity of the deconvoluted mass peaks was assigned initially based on the expected mass of the purified proteins, and subsequently based on the observed mass shifts between peaks in the deconvoluted mass spectra. The lower molecular weight CerS6 peak corresponds to loss of the initiator methionine (theoretical mass shift of -131.20 Da), acetylation of the new N-terminus (theoretical: +42.04 Da) and addition of an N-linked GlcNAc (theoretical: +203.19 Da). However, the major glycosylation species observed contains the complete core N-linked glycan (theoretical: +1217.05 Da), as expected for protein expressed in Expi293F GnTI-cells. In addition, we observed +237.47 and +238.51 Da mass shifts relatively to these two glycosylated species, respectively, which we interpreted as palmitoylation at a single site (theoretical: +238.41 Da), corresponding to the covalent acyl-imidazole intermediate observed in the cryo-EM structure. One additional modification was observed (∼ +264 Da), but its identity could not be assigned. This unknown modification occurs at a distinct site to that of the palmitoylation, as both modifications can occur simultaneously in the same protein molecule (Extended Data Fig. 1f,g), thus excluding the possibility of this unknown modification occurring in the active site.

To monitor the reaction of the covalent intermediate species with the second substrates, prior to denaturing intact protein mass spectrometry analysis the protein (2 mg mL^-1^) was incubated with 200 µM sphinganine (Avanti Polar Lipids), 200 µM fumonisin B_1_ (Sigma-Aldrich), or 600 µM FTY720 (Sigma-Aldrich) for 1h 30 min at 37 °C. All intact mass experiments were conducted at least twice using distinct biological samples. Replicate deconvoluted mass spectra are shown in Extended Data Fig. 5.

### Product detection by liquid chromatography high-resolution mass spectrometry (LC-HRMS)

Product detection by LC-HRMS was conducted on a nanoElute LC system in line with a TIMS TOF Pro 2 mass spectrometer (Bruker). The reactions were set up as described for the denaturing intact protein MS experiments, diluted 1:40 (v/v) in 30 % methanol in 0.1 % formic acid, and 1 µL of each sample was injected onto an IonOpticks C18 nano UHPLC column (1.6 μm particle size; 0.075 x 250 mm).

The flow rate was set to 0.5 µL min^-1^, and the solvent system consisted of 0.1 % Optima LC– MS grade formic acid (Fisher Chemical) in high-performance liquid chromatography electrochemical grade water (Fisher Chemical) (solvent A) and 0.1 % formic acid in Optima LC–MS grade methanol (Fisher Chemical) (solvent B). Initial conditions were 60 % solvent B, and a linear gradient from 60 % to 95 % solvent B was applied over 17.8 min to elute the samples. This was then followed by a final 2.2 min isocratic elution with 95 % solvent B, before the system was re-equilibrated between samples for 5 min with 60 % solvent B.

The mass spectrometer was operated in positive ion mode with a capillary voltage of 1,600 V, and the drying gas was supplied at 180 °C with a flow rate of 3 L min^−^1. Additional parameters were: deflection 1 delta of 70 V, funnel 1 RF of 350 Vpp, funnel 2 RF of 600 Vpp, multipole RF of 500 Vpp. Data were acquired between 150-2200 *m/z*, and analyzed using the Bruker Compass DataAnalysis 5.3.556 software. The extracted ion chromatograms are presented in Fig. 3c and Fig. 4d, and correspond to the theoretical [M+H]^+^ ions (tolerance: ± 0.005 *m/z*) of the reaction products N-palmitoyl dihydrosphingosine (C16:0 ceramide) (theoretical *m/z*: 540.5350), N-palmitoyl FB_1_ (theoretical *m/z*: 960.6254), and N-palmitoyl FTY720 (theoretical *m/z*: 546.4881).

### Liquid chromatography electrospray ionization tandem mass spectrometry (LC-ESI-MS/MS) characterization of N-palmitoyl FTY720

To structurally characterize the proposed N-palmitoyl FTY720 reaction product, purified protein was incubated with 600 μM FTY720 as before, and the reaction mixture was initially analyzed by LC-ESI-MS on an Agilent 1290 Infinity LC System in line with an Agilent 6530 Accurate-Mass quadrupole time-of-flight mass spectrometer (Agilent Technologies) as described above. This enabled the identification of the putative [M+H]^+^ ion of the N-palmitoyl FTY720 product (observed *m/z*: 546.4849; theoretical *m/z*: 546.4881). Its product ion spectrum was then obtained by LC-ESI-MS/MS. For this purpose, the mass spectrometer was operated in positive electrospray ionization (ESI+), 4 GHz mode. MS parameters were capillary voltage of 4,000 V, fragmentor voltage of 175 V. Data were acquired between 100-1700 *m/z*. The targeted parent ion (546.4881 *m/z*; retention time = 8.713 min) was fragmented using a collision energy of 14 V.

### Dihydroceramide synthase activity measurements

For activity assays, wild-type or mutant CerS6 proteins were over-expressed in Expi293F cells, and membranes were prepared as follows. Cell pellet from 0.5 L of culture was thawed in PBS, lysed using an Emulsiflex C5 homogeniser (Avestin), and cell debris was removed by centrifugation. Membranes were subsequently isolated by ultracentrifugation at 160,000 x g for 1h 30 min, resuspended in Assay Buffer (20 mM HEPES pH 7.5, 25 mM KCl, 1 mM MgSO_4_, 0.1 % (v/v) glycerol), flash-frozen in liquid N_2_ and stored at -80 °C.

On the day of the assay, membranes were thawed, diluted to 0.25 mg mL^-1^, and 20 µL of diluted membranes were dispensed per well on flat bottom polystyrene 384-well Lumitrac plates (Greiner Bio One). Then, 50 µM sphinganine (2.5 µL 500 µM sphinganine in Assay Buffer containing 10% ethanol) and 50 µM palmitoyl-CoA (Sigma-Aldrich) (2.5 µL 500 µM palmitoyl-CoA in Assay Buffer) were added to each well for a final reaction volume of 25 µL. For untreated controls, 5 µL of Assay Buffer were added instead of the substrates. The plates were then incubated for 1h at room temperature, and the reaction was terminated by the addition of 40 µL butanol spiked with N-palmitoyl(d9) dihydrosphingosine (Avanti Polar Lipids) to yield a final concentration of 5 µM N-palmitoyl(d9) dihydrosphingosine as an internal standard. The plates were then shaken at 1,800 rpm for 2 min, centrifuged at 1,000 x g for 30 s, and 40 µL of the organic (upper) phase were transferred into 384-well polypropylene deep well plates (Greiner Bio One) and diluted with 40 µL butanol.

The analytical sample handling was performed by a rapid-injecting RapidFire autosampler system (Agilent, Waldbronn, Germany) coupled to a triple quadrupole mass spectrometer (Triple Quad 6500, AB Sciex Germany GmbH, Darmstadt, Germany) as previously described^73^, with the following modifications. Liquid sample is aspirated by a vacuum pump into a 10 µL sample loop for 6000 ms and subsequently flushed for 3000 ms onto a C4 cartridge (Agilent) with the aqueous mobile phase (99.5% water, 0.49% acetic Acid, 0.01% Trifluoroacetic acid, flow rate 1.5 mL min^-1^). The MRM transition for C16:0 dihydroceramide is 540.5 → 266.3 *m/z* (declustering potential 130V, collision energy 38 V) and for the internal standard C16:0 (d9)dihydroceramide is 549.5 → 266.3 *m/z* (declustering potential 130V, collision energy 38 V). The mass spectrometer was operated in positive ionization mode (curtain gas 35 a.u., collision gas medium, ion spray voltage 4200 V, temperature 550 °C, ion source gas 1 65 a.u., ion source gas 2 80 a.u.). MS data processing was performed in GMSU (Alpharetta, GA, USA), and peak area ratios between C16:0 dihydroceramide and internal standard peaks were calculated.

Expression of each mutant in the membrane samples was evaluated through western blotting using 10 ng mL^-1^ Strep-Tactin conjugated with horseradish peroxidase (IBA Lifesciences) and using SYPRO Ruby (Thermo Fisher Scientific) staining as a loading control (Source Data Fig. 1). Band intensity was quantified using the GeneTools software (Syngene) and activity was normalized to protein expression.

### Molecular dynamics simulations

Atomistic molecular dynamics simulations were performed using the Desmond software package (D. E. Shaw Research, New York, NY) within the Maestro software suite (Schrödinger). The simulation setup and analysis were carried out as follows: For each of the two states, a CerS6 monomer was embedded in a lipid bilayer consisting of 99 POPC molecules after undergoing the protein preparation step as implemented in Maestro. The dimensions of the simulation cell were approximately 90x70x70 Å, with a minimum distance of 10 Å between the protein and the cell boundaries. The systems contained 8,855 (covalent intermediate bound state) or 9,368 (N-acyl FB_1_ bound state) water molecules and correspondingly 8 or 6 chloride counterions to maintain overall charge neutrality. The simulations were performed using the OPLS4 force field^74^ for the protein and lipid molecules, and the SPC water model for the water molecules. The system was set up using the Maestro software suite (Schrödinger). The simulations were run in Desmond using an NPT ensemble at a temperature of 300K. A timestep of 2 fs was employed for the integration of equations of motion. Four independent simulations were performed, each with a duration of 100 ns, resulting in a total simulation time of 400 ns. Snapshots of the system were recorded every 100 ps for further analysis.

## Figures

Figures depicting molecular models were generated using PyMOL^75^ and ChimeraX^76^.

## Data availability

The cryo-EM maps have been deposited in the Electron Microscopy Data Bank (EMDB) under accession codes EMD-18770 (covalent acyl-enzyme intermediate state) and EMD-18771 (N-acyl FB_1_-bound state). The atomic models have been deposited in the Protein Data Bank under accession codes 8QZ6 (covalent acyl-enzyme intermediate state) and 8QZ7 (N-acyl FB_1_-bound state).

## Acknowledgements

T.C.P. was supported by a Wellcome PhD studentship (102164/B/15/Z). Initial work on this project was funded by the Structural Genomics Consortium, a registered charity (number 1097737) that received funds from AbbVie, Bayer, Boehringer Ingelheim, the Canada Foundation for Innovation, Genome Canada, Janssen, Merck, Novartis, the Ontario Ministry of Economic Development and Innovation, Pfizer, and Takeda, as well as the Wellcome Trust (106169/Z/14/Z). D.B.S., A.C.W.P. and G.C. were supported by the Innovative Medicines Initiative 2 Joint Undertaking (JU) under grant agreement No 875510. The JU receives support from the European Union’s Horizon 2020 research and innovation program and EFPIA and Ontario Institute for Cancer Research, Royal Institution for the Advancement of Learning McGill University, Kungliga Tekniska Hoegskolan, Diamond Light Source Limited. Electron microscopy was provided through the Oxford Particle Imaging Centre (OPIC), an Instruct-ERIC centre (funded by Wellcome Trust JIF award [060208/Z/00/Z] and equipment grant [093305/Z/10/Z]) and the Midlands Regional CryoEM Facility at the Leicester Institute of Structural and Chemical Biology (LISCB), major funding from MRC (MC_PC_17136). We thank Christos S. Savva (LISCB) for technical assistance with cryo-EM data collection and William Greenland (Agilent) for technical advice on tandem mass spectrometry. We thank Wyatt Yue and Christian Siebold for helpful discussions.

## Author contributions

T.C.P. expressed and purified protein samples, prepared proteoliposome samples for alpaca immunization and membranes for mutant activity testing, carried out biophysical characterizations including bio-layer interferometry experiments, designed and generated CerS6 mutants by site-directed mutagenesis, built and refined the atomic models, and performed intact protein MS, product identification by high-resolution MS and small-molecule structural characterization by LC-ESI-MS/MS. T.C.P. and G.C. prepared cryo-EM grids. T.C.P. and A.C.W.P. collected and processed the cryo-EM data. C.S.T. carried out the molecular dynamics simulations and analysis. A.Q. was involved in the early stages of the project, including initial screening of expression and purification conditions. R.C. provided access to mass spectrometry instruments for intact protein and small-molecule analysis, and assisted with small-molecule MS experiments. S.Š. conducted alpaca immunizations, nanobody library generation, panning and identification of unique nanobody sequences. M.T. performed the CerS6 mutant activity assays, supervised by S.T.. T.C.P. and D.B.S. wrote the original draft of the manuscript. All authors participated in the discussion and manuscript editing. A.P., E.P.C., G.S. and D.B.S. supervised the research.

## Competing interests

The authors declare no competing interests. C.S.T., M.T., S.T., A.P., and G.S. are employees of Boehringer Ingelheim Pharma GmbH & Co. KG.

## Notes

### Competing Interest Statement

The authors have declared no competing interest.

### Summary of Updates

Supplementary Material uploaded

## References

1 Summers, S. A., Chaurasia, B. & Holland, W. L. Metabolic Messengers: ceramides. Nature Metabolism 1, 1051–1058, doi:10.1038/s42255-019-0134-8 (2019).

2 Turpin, S. M. et al. Obesity-induced CerS6-dependent C16:0 ceramide production promotes weight gain and glucose intolerance. Cell Metab 20, 678–686, doi:10.1016/j.cmet.2014.08.002 (2014).

3 Apostolopoulou, M. et al. Specific Hepatic Sphingolipids Relate to Insulin Resistance, Oxidative Stress, and Inflammation in Nonalcoholic Steatohepatitis. Diabetes Care 41, 1235–1243, doi:10.2337/dc17-1318 (2018).

4 Hajduch, E., Lachkar, F., Ferre, P. & Foufelle, F. Roles of Ceramides in Non-Alcoholic Fatty Liver Disease. J Clin Med 10, 792, doi:10.3390/jcm10040792 (2021).

5 Hammerschmidt, P. & Brüning, J. C. Contribution of specific ceramides to obesity-associated metabolic diseases. Cell Mol Life Sci 79, 395, doi:10.1007/s00018-022-04401-3 (2022).

6 Neeland, I. J. et al. Relation of plasma ceramides to visceral adiposity, insulin resistance and the development of type 2 diabetes mellitus: the Dallas Heart Study. Diabetologia 61, 2570–2579, doi:10.1007/s00125-018-4720-1 (2018).

7 Błachnio-Zabielska, A. U. et al. Increased bioactive lipids content in human subcutaneous and epicardial fat tissue correlates with insulin resistance. Lipids 47, 1131–1141, doi:10.1007/s11745-012-3722-x (2012).

8 Luukkonen, P. K. et al. Hepatic ceramides dissociate steatosis and insulin resistance in patients with non-alcoholic fatty liver disease. J Hepatol 64, 1167–1175, doi:10.1016/j.jhep.2016.01.002 (2016).

9 Levy, M. & Futerman, A. H. Mammalian ceramide synthases. IUBMB Life 62, 347–356, doi:10.1002/iub.319 (2010).

10 Ternes, P., Franke, S., Zähringer, U., Sperling, P. & Heinz, E. Identification and characterization of a sphingolipid delta 4-desaturase family. J Biol Chem 277, 25512–25518, doi:10.1074/jbc.M202947200 (2002).

11 Kitatani, K., Idkowiak-Baldys, J. & Hannun, Y. A. The sphingolipid salvage pathway in ceramide metabolism and signaling. Cell Signal 20, 1010–1018, doi:10.1016/j.cellsig.2007.12.006 (2008).

12 Raichur, S. et al. The role of C16:0 ceramide in the development of obesity and type 2 diabetes: CerS6 inhibition as a novel therapeutic approach. Mol Metab 21, 36–50, doi:10.1016/j.molmet.2018.12.008 (2019).

13 Raichur, S. Ceramide Synthases Are Attractive Drug Targets for Treating Metabolic Diseases. Front Endocrinol 11, 483, doi:10.3389/fendo.2020.00483 (2020).

14 Hammerschmidt, P. et al. CerS6-Derived Sphingolipids Interact with Mff and Promote Mitochondrial Fragmentation in Obesity. Cell 177, 1536–1552.e1523, doi:10.1016/j.cell.2019.05.008 (2019).

15 Mizutani, Y., Kihara, A. & Igarashi, Y. Mammalian Lass6 and its related family members regulate synthesis of specific ceramides. Biochemical Journal 390, 263–271, doi:10.1042/bj20050291 (2005).

16 Winter, E. & Ponting, C. P. TRAM, LAG1 and CLN8: members of a novel family of lipid-sensing domains? Trends Biochem Sci 27, 381–383, doi:10.1016/s0968-0004(02)02154-0 (2002).

17 Spassieva, S. et al. Necessary role for the Lag1p motif in (dihydro)ceramide synthase activity. J Biol Chem 281, 33931–33938, doi:10.1074/jbc.M608092200 (2006).

18 Kageyama-Yahara, N. & Riezman, H. Transmembrane topology of ceramide synthase in yeast. Biochem J 398, 585–593, doi:10.1042/bj20060697 (2006).

19 Venkataraman, K. & Futerman, A. H. Do longevity assurance genes containing Hox domains regulate cell development via ceramide synthesis? FEBS Lett 528, 3–4, doi:10.1016/s0014-5793(02)03248-9 (2002).

20 Mesika, A., Ben-Dor, S., Laviad, E. L. & Futerman, A. H. A new functional motif in Hox domain-containing ceramide synthases: identification of a novel region flanking the Hox and TLC domains essential for activity. J Biol Chem 282, 27366–27373, doi:10.1074/jbc.M703487200 (2007).

21 Sassa, T., Hirayama, T. & Kihara, A. Enzyme Activities of the Ceramide Synthases CERS2-6 Are Regulated by Phosphorylation in the C-terminal Region. J Biol Chem 291, 7477–7487, doi:10.1074/jbc.M115.695858 (2016).

22 Tidhar, R. et al. Eleven residues determine the acyl chain specificity of ceramide synthases. J Biol Chem 293, 9912–9921, doi:10.1074/jbc.RA118.001936 (2018).

23 Laviad, E. L., Kelly, S., Merrill, A. H., Jr. & Futerman, A. H. Modulation of ceramide synthase activity via dimerization. J Biol Chem 287, 21025–21033, doi:10.1074/jbc.M112.363580 (2012).

24 Jensen, S. A. et al. Bcl2L13 is a ceramide synthase inhibitor in glioblastoma. Proc Natl Acad Sci U S A 111, 5682–5687, doi:10.1073/pnas.1316700111 (2014).

25 Kim, J. L. et al. Fatty acid transport protein 2 interacts with ceramide synthase 2 to promote ceramide synthesis. Journal of Biological Chemistry 298, 101735, doi:10.1016/j.jbc.2022.101735 (2022).

26 Cingolani, F., Futerman, A. H. & Casas, J. Ceramide synthases in biomedical research. Chem Phys Lipids 197, 25–32, doi:10.1016/j.chemphyslip.2015.07.026 (2016).

27 Marasas, W. F. et al. Fumonisins disrupt sphingolipid metabolism, folate transport, and neural tube development in embryo culture and in vivo: a potential risk factor for human neural tube defects among populations consuming fumonisin-contaminated maize. J Nutr 134, 711–716, doi:10.1093/jn/134.4.711 (2004).

28 Riley, R. T. & Merrill, A. H., Jr. Ceramide synthase inhibition by fumonisins: a perfect storm of perturbed sphingolipid metabolism, signaling, and disease. J Lipid Res 60, 1183–1189, doi:10.1194/jlr.S093815 (2019).

29 Riley, R. T. et al. Fumonisins (addendum). 415–573 (World Health Organization, Geneva, Switzerland, 2018).

30 Merrill, A. H., Jr., van Echten, G., Wang, E. & Sandhoff, K. Fumonisin B_1_ inhibits sphingosine (sphinganine) N-acyltransferase and de novo sphingolipid biosynthesis in cultured neurons in situ. J Biol Chem 268, 27299–27306 (1993).

31 Berdyshev, E. V. et al. FTY720 Inhibits Ceramide Synthases and Up-regulates Dihydrosphingosine 1-Phosphate Formation in Human Lung Endothelial Cells. Journal of Biological Chemistry 284, 5467–5477, doi:10.1074/jbc.m805186200 (2009).

32 Lahiri, S. et al. Ceramide synthesis is modulated by the sphingosine analog FTY720 via a mixture of uncompetitive and noncompetitive inhibition in an Acyl-CoA chain length-dependent manner. J Biol Chem 284, 16090–16098, doi:10.1074/jbc.M807438200 (2009).

33 Turner, N. et al. A selective inhibitor of ceramide synthase 1 reveals a novel role in fat metabolism. Nature Communications 9, 3165, doi:10.1038/s41467-018-05613-7 (2018).

34 Nie, L. et al. The structural basis of fatty acid elongation by the ELOVL elongases. Nature Structural & Molecular Biology 28, 512–520, doi:10.1038/s41594-021-00605-6 (2021).

35 Niu, Y. et al. Analysis of the mechanosensor channel functionality of TACAN. eLife 10, e71188 , citation = eLife 72021;71110 e71188, doi:10.7554/eLife.71188 (2021).

36 Ke, M. et al. Cryo-EM structures of human TMEM120A and TMEM120B. Cell Discov 7, 77, doi:10.1038/s41421-021-00319-5 (2021).

37 Holm, L., Laiho, A., Törönen, P. & Salgado, M. DALI shines a light on remote homologs: One hundred discoveries. Protein Sci 32, e4519, doi:10.1002/pro.4519 (2023).

38 Ikeda, M. et al. Characterization of four mammalian 3-hydroxyacyl-CoA dehydratases involved in very long-chain fatty acid synthesis. FEBS Letters 582, 2435–2440, doi:10.1016/j.febslet.2008.06.007 (2008).

39 Cleland, W. W. The kinetics of enzyme-catalyzed reactions with two or more substrates or products. I. Nomenclature and rate equations. Biochim Biophys Acta 67, 104–137, doi:10.1016/0006-3002(63)91800-6 (1963).

40 Zollinger, M. et al. Absorption and disposition of the sphingosine 1-phosphate receptor modulator fingolimod (FTY720) in healthy volunteers: a case of xenobiotic biotransformation following endogenous metabolic pathways. Drug Metab Dispos 39, 199–207, doi:10.1124/dmd.110.035907 (2011).

41 Wedekind, J. E., Frey, P. A. & Rayment, I. The structure of nucleotidylated histidine-166 of galactose-1-phosphate uridylyltransferase provides insight into phosphoryl group transfer. Biochemistry 35, 11560–11569, doi:10.1021/bi9612677 (1996).

42 Zhou, X. et al. Kinetic Mechanism of Human Histidine Triad Nucleotide Binding Protein 1. Biochemistry 52, 3588–3600, doi:10.1021/bi301616c (2013).

43 Trent, M. S., Worsham, L. M. S. & Ernst-Fonberg, M. L. HlyC, the Internal Protein Acyltransferase That Activates Hemolysin Toxin: Roles of Various Conserved Residues in Enzymatic Activity As Probed by Site-Directed Mutagenesis. Biochemistry 38, 9541–9548, doi:10.1021/bi9905617 (1999).

44 Dodds, A. W., Ren, X. D., Willis, A. C. & Law, S. K. The reaction mechanism of the internal thioester in the human complement component C4. Nature 379, 177–179, doi:10.1038/379177a0 (1996).

45 Gadjeva, M. et al. The covalent binding reaction of complement component C3. J Immunol 161, 985–990 (1998).

46 Selvy, P. E., Lavieri, R. R., Lindsley, C. W. & Brown, H. A. Phospholipase D: enzymology, functionality, and chemical modulation. Chem Rev 111, 6064–6119, doi:10.1021/cr200296t (2011).

47 Vanni, N. et al. Impairment of ceramide synthesis causes a novel progressive myoclonus epilepsy. Ann Neurol 76, 206–212, doi:10.1002/ana.24170 (2014).

48 Harrer, H., Laviad, E. L., Humpf, H. U. & Futerman, A. H. Identification of N-acyl-fumonisin B_1_ as new cytotoxic metabolites of fumonisin mycotoxins. Mol Nutr Food Res 57, 516–522, doi:10.1002/mnfr.201200465 (2013).

49 Harrer, H., Humpf, H. U. & Voss, K. A. In vivo formation of N-acyl-fumonisin B_1_. Mycotoxin Res 31, 33–40, doi:10.1007/s12550-014-0211-5 (2015).

50 Zelnik, I. D. et al. Computational design and molecular dynamics simulations suggest the mode of substrate binding in ceramide synthases. Nat Commun 14, 2330, doi:10.1038/s41467-023-38047-x (2023).

51 Youssefian, L. et al. Autosomal recessive congenital ichthyosis: CERS3 mutations identified by a next generation sequencing panel targeting ichthyosis genes. Eur J Hum Genet 25, 1282–1285, doi:10.1038/ejhg.2017.137 (2017).

52 Eckl, K. M. et al. Impaired epidermal ceramide synthesis causes autosomal recessive congenital ichthyosis and reveals the importance of ceramide acyl chain length. J Invest Dermatol 133, 2202–2211, doi:10.1038/jid.2013.153 (2013).

53 Radner, F. P. et al. Mutations in CERS3 cause autosomal recessive congenital ichthyosis in humans. PLoS Genet 9, e1003536, doi:10.1371/journal.pgen.1003536 (2013).

54 Dougherty, D. A. The cation-π interaction. Acc Chem Res 46, 885–893, doi:10.1021/ar300265y (2013).

55 Merrill, A. H., Jr., Wang, E., Gilchrist, D. G. & Riley, R. T. Fumonisins and other inhibitors of de novo sphingolipid biosynthesis. Adv Lipid Res 26, 215–234 (1993).

56 Xie, T. et al. Structure and mechanism of a eukaryotic ceramide synthase complex. Embo j, e114889, doi:10.15252/embj.2023114889 (2023).

57 Tidhar, R., Sims, K., Rosenfeld-Gur, E., Shaw, W. & Futerman, A. H. A rapid ceramide synthase activity using NBD-sphinganine and solid phase extraction. J Lipid Res 56, 193–199, doi:10.1194/jlr.D052001 (2015).

58 Geertsma, E. R., Nik Mahmood, N. A., Schuurman-Wolters, G. K. & Poolman, B. Membrane reconstitution of ABC transporters and assays of translocator function. Nat Protoc 3, 256–266, doi:10.1038/nprot.2007.519 (2008).

59 Hutter, C. A. J. et al. The extracellular gate shapes the energy profile of an ABC exporter. Nature Communications 10, 2260, doi:10.1038/s41467-019-09892-6 (2019).

60 Pardon, E. et al. A general protocol for the generation of Nanobodies for structural biology.Nature Protocols 9, 674–693, doi:10.1038/nprot.2014.039 (2014).

61 Zheng, S. Q. et al. MotionCor2: anisotropic correction of beam-induced motion for improved cryo-electron microscopy. Nat Methods 14, 331–332, doi:10.1038/nmeth.4193 (2017).

62 Punjani, A., Rubinstein, J. L., Fleet, D. J. & Brubaker, M. A. cryoSPARC: algorithms for rapid unsupervised cryo-EM structure determination. Nat Methods 14, 290–296, doi:10.1038/nmeth.4169 (2017).

63 Punjani, A., Zhang, H. & Fleet, D. J. Non-uniform refinement: adaptive regularization improves single-particle cryo-EM reconstruction. Nat Methods 17, 1214–1221, doi:10.1038/s41592-020-00990-8 (2020).

64 Asarnow, D., Palovcak, E. & Cheng, Y. UCSF pyem v0.5, <10.5281/zenodo.3576630}> (2019).

65 Zivanov, J. et al. New tools for automated high-resolution cryo-EM structure determination in RELION-3. Elife 7, doi:10.7554/eLife.42166 (2018).

66 Jumper, J. et al. Highly accurate protein structure prediction with AlphaFold. Nature 596, 583–589, doi:10.1038/s41586-021-03819-2 (2021).

67 Kelley, L. A., Mezulis, S., Yates, C. M., Wass, M. N. & Sternberg, M. J. The Phyre2 web portal for protein modeling, prediction and analysis. Nat Protoc 10, 845–858, doi:10.1038/nprot.2015.053 (2015).

68 Emsley, P., Lohkamp, B., Scott, W. G. & Cowtan, K. Features and development of Coot. *Acta crystallographica*. Section D, Biological crystallography 66, 486–501, doi:10.1107/S0907444910007493 (2010).

69 Afonine, P. V. et al. Real-space refinement in PHENIX for cryo-EM and crystallography. Acta Crystallogr D Struct Biol 74, 531–544, doi:10.1107/s2059798318006551 (2018).

70 Long, F. et al. AceDRG: a stereochemical description generator for ligands. Acta Crystallogr D Struct Biol 73, 112–122, doi:10.1107/s2059798317000067 (2017).

71 Yamashita, K., Palmer, C. M., Burnley, T. & Murshudov, G. N. Cryo-EM single-particle structure refinement and map calculation using Servalcat. Acta Crystallogr D Struct Biol 77, 1282–1291, doi:10.1107/s2059798321009475 (2021).

72 Burnley, T., Palmer, C. M. & Winn, M. Recent developments in the CCP-EM software suite. Acta Crystallogr D Struct Biol 73, 469–477, doi:10.1107/s2059798317007859 (2017).

73 Tautermann, C. S. et al. Allosteric Activation of Striatal-Enriched Protein Tyrosine Phosphatase (STEP, PTPN5) by a Fragment-like Molecule. J Med Chem 62, 306-316, doi:10.1021/acs.jmedchem.8b00857 (2019).

74 Lu, C. et al. OPLS4: Improving Force Field Accuracy on Challenging Regimes of Chemical Space. J Chem Theory Comput 17, 4291–4300, doi:10.1021/acs.jctc.1c00302 (2021).

75 Schrodinger, LLC. The PyMOL Molecular Graphics System, Version 2.*1* (2015).

76 Pettersen, E. F. et al. UCSF ChimeraX: Structure visualization for researchers, educators, and developers. Protein Sci 30, 70–82, doi:10.1002/pro.3943 (2021).

