## Extended Data for "Structural basis of the mechanism and inhibition of a human ceramide synthase"

**Extended Data Table 1: Cryo-EM data collection, processing, refinement and model validation statistics**

| | CerS6<br>covalent intermediate state | CerS6<br>N-palm $FB_1$-bound state |
|---|---|---|
| EMDB ID | EMD-18770 | EMD-18771 |
| PDB ID | 8QZ6 | 8QZ7 |
| **Data collection and processing** | | |
| Microscope | Krios | Krios |
| Detector | K3 | K3 |
| Voltage (kV) | 300 | 300 |
| Nominal magnification | 130,000 | 130,000 |
| Pixel size (Å $px^{-1}$) | 0.656 | 0.656 |
| Total dose ($e^{-}$ $A^{-2}$) | 56.3 | 56.3 |
| Defocus range (µm) | -0.8 to -2.4 | -0.8 to -2.4 |
| Movies collected (used) | 14,309 (14,196) | 18,386 (16,647) |
| Initial particles | 3,400,118 | 3,998,935 |
| Particles post 2D | 1,261,928 | 1,353,329 |
| Final particles | 93,680 | 154,239 |
| Symmetry imposed | C2 | C2 |
| Map resolution (Å) GSFSC = 0.143 | 3.22 | 2.95 |
| **Refinement** | | |
| Map sharpening B-factor (Å$^2$) | -132.5 | -130.7 |
| Map CC (phenix CC mask) | 0.83 | 0.85 |
| Map-to-model resolution (FSC = 0.5) | 3.38 | 3.22 |
| **Model building and validation** | | |
| **Model composition** | | |
| Non-hydrogen protein atoms | 7,226 | 7,366 |
| Protein residues | 906 | 914 |
| Ligands | | |
| Palmitoyl chain | 2 | / |
| N-palmitoyl $FB_1$ | / | 2 |
| POPC | 2 | 2 |
| GlcNAc | 2 | 2 |
| **RMSD from ideal** | | |
| Bond length (Å) | 0.002 | 0.003 |
| Bond angles (º) | 0.393 | 0.458 |
| **Validation** | | |
| Molprobity score | 1.15 | 1.20 |
| Clashscore | 3.62 | 3.17 |
| Rotamer outliers (%) | 0.29 | 0.87 |
| Ramachandran favoured (%) | 98.00 | 97.57 |
| Ramachandran allowed (%) | 2.00 | 2.43 |
| Ramachandran outliers (%) | 0.00 | 0.00 |

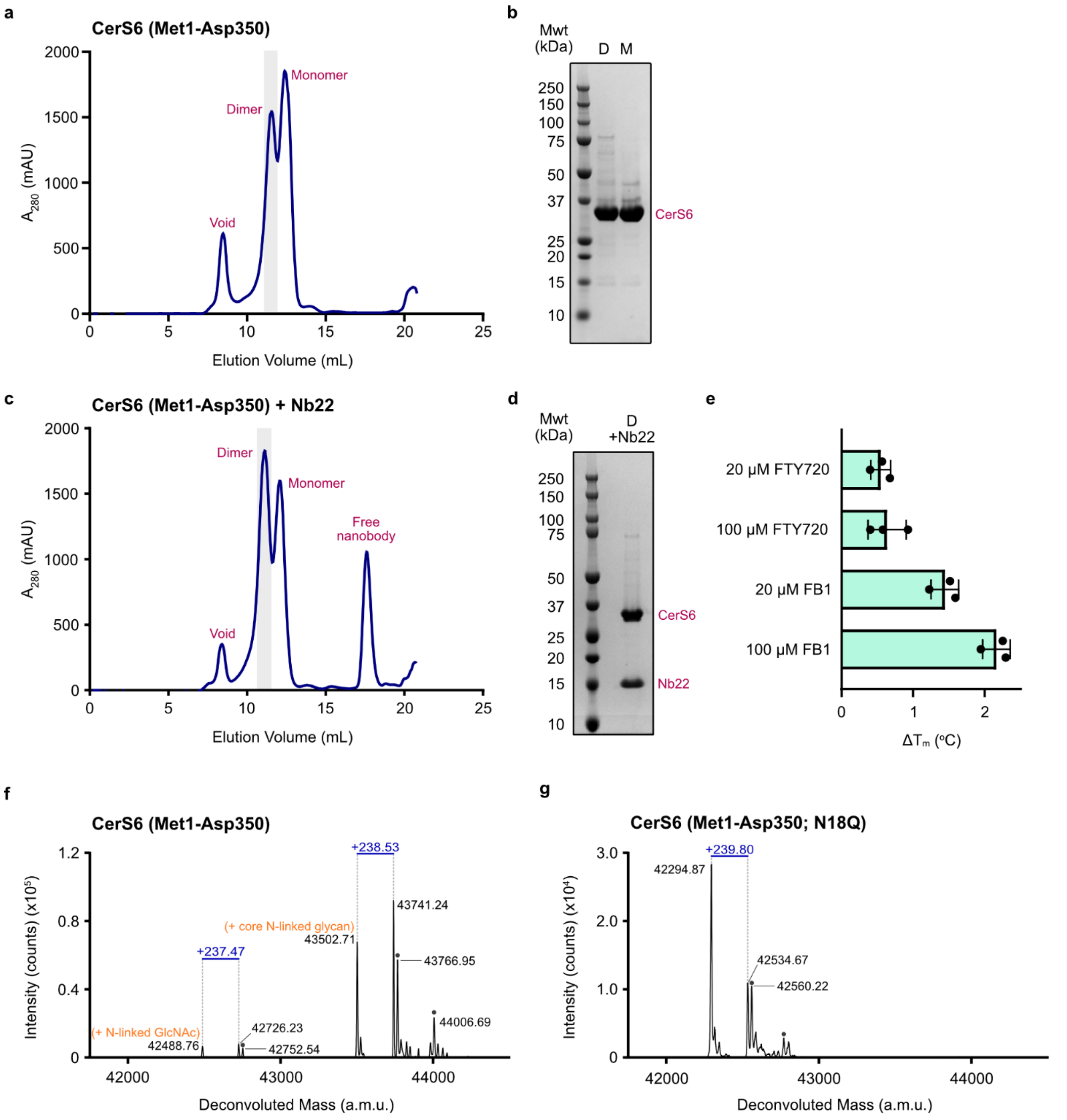


**Extended Data Figure 1. Properties of purified CerS6. a**, Elution profile of CerS6 (Met1-Asp350). The collected dimer peak is shaded in gray. **b**, SDS-PAGE analysis of the purified dimeric (D) and monomeric (M) SEC peak fractions. **c**, Elution profile of CerS6 (Met1-Asp350) in complex with nanobody 22. The putative dimer peak, shaded in gray, was pooled and concentrated for single particle Cryo-EM. **d**, SDS-PAGE analysis of the SEC-purified CerS6 dimer and nanobody 22 complex. **e,** NanoDSF screening of CerS inhibitors. Purified CerS6 (Met1-Asp350) was incubated with 20 µM or 100 µM of each compound for 1h on ice prior to thermal unfolding. **f-g**, Denaturing intact protein MS analysis of purified CerS6. **f**, Deconvoluted mass spectra of wild-type CerS6 (Met1-Asp350). The expected mass of the untagged, unmodified, truncated enzyme based on the sequence is

42,373.53 Da. The observed lower mass peak (42,488.76 Da) corresponds to loss of the initiator methionine (-131.20 Da), acetylation of the new N-terminus (+42.04 Da), and addition of an N-linked GlcNAc (+203.19 Da). An additional, higher intensity, deconvoluted mass peak (43,502.71 Da) corresponds to the addition of a core N-linked glycan (theoretical +1217.09 Da) in place of simply the N-linked GlcNAc. Mass shifts corresponding to the mass of a palmitoyl group (theoretical +238.41 Da) are labelled in blue. Additional deconvoluted mass peaks corresponding to the addition of an unknown modification of approximately +264 Da on top of the glycosylation or glycosylation + palmitoylation peaks are labelled with gray dots. **g**, Deconvoluted mass spectra obtained for the purified CerS6 N18Q mutant which removes the only identified glycosylation site. The expected mass of the untagged, unmodified, truncated CerS6 N18Q enzyme based on the sequence is 42,387.56 Da. The observed lower mass peak (42,294.87 Da) corresponds to loss of the initiator methionine followed by acetylation of the new N-terminus. The +239.80 Da mass shift, corresponding to palmitoylated protein, is labelled in blue.

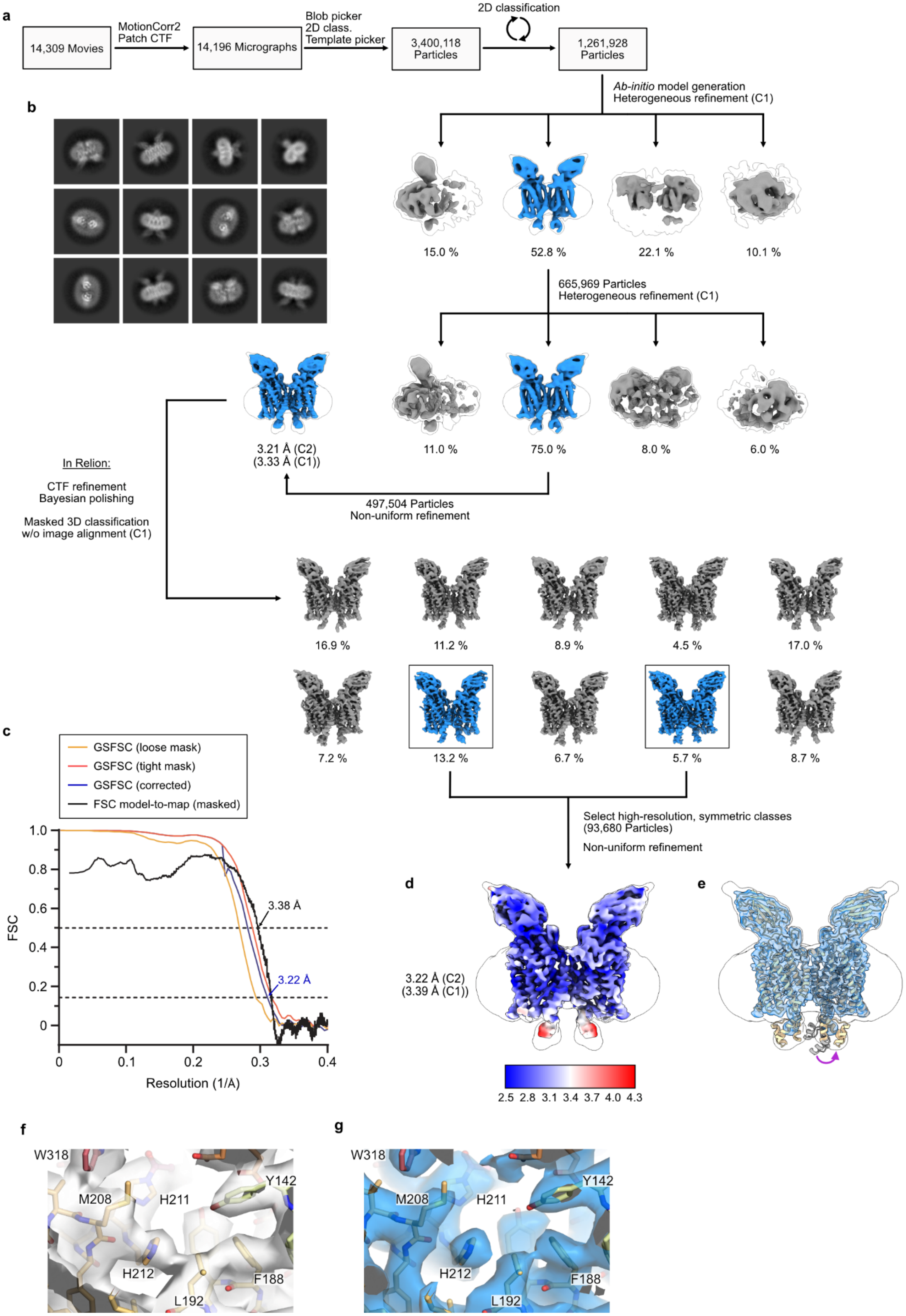

a
14,309 Movies
MotionCorr2
Patch CTF
14,196 Micrographs
Blob picker
2D class.
Template picker
3,400,118
Particles
2D classification
1,261,928
Particles
Ab-initio model generation
Heterogeneous refinement (C1)
15.0 %
52.8 %
22.1 %
10.1 %
665,969 Particles
Heterogeneous refinement (C1)
11.0 %
75.0 %
8.0 %
6.0 %
3.21 Å (C2)
(3.33 Å (C1))
497,504 Particles
Non-uniform refinement
b
In Relion:
CTF refinement
Bayesian polishing
Masked 3D classification
w/o image alignment (C1)
16.9 %
11.2 %
8.9 %
4.5 %
17.0 %
7.2 %
13.2 %
6.7 %
5.7 %
8.7 %
Select high-resolution, symmetric classes
(93,680 Particles)
Non-uniform refinement
c
GSFSC (loose mask)
GSFSC (tight mask)
GSFSC (corrected)
FSC model-to-map (masked)
3.38 Å
3.22 Å
FSC
Resolution (1/Å)
1.0
0.8
0.6
0.4
0.2
0
0.1
0.2
0.3
0.4
d
3.22 Å (C2)
(3.39 Å (C1))
2.5 2.8 3.1 3.4 3.7 4.0 4.3
e
f
W318
M208
H211
Y142
H212
F188
L192
g
W318
M208
H211
Y142
H212
F188
L192

**Extended Data Figure 2. Cryo-EM data processing: covalent intermediate dataset. a**, Cryo-EM data processing flowchart. **b**, 2D classes obtained prior to 3D classification. **c**, Fourier Shell Correlation (FSC) plots, indicating overall map resolution (GSFSC = 0.143) and a model-to-map FSC curve. **d,** Final 3D reconstruction, colored by local resolution. **e,** Overlay of the structural model (tan) and the AlphaFold2 monomer prediction (gray). The Cryo-EM maps are shown as blue (unsharpened map) or outline (blurred map; $B_{blur}$ = 200 Å$^2$) surfaces, revealing that density corresponding to the Hox-like domains is clearly visible in the blurred maps. A purple arrow indicates the manual adjustment of the position of the Hox-like domain upon docking of its AlphaFold2 prediction into the experimental blurred maps. **f-g,** Cryo-EM map **(f)** before and **(g)** after CTF refinement, Bayesian polishing and 3D classification in Relion, revealing an improvement in map quality in the final map despite no improvement in nominal resolution.

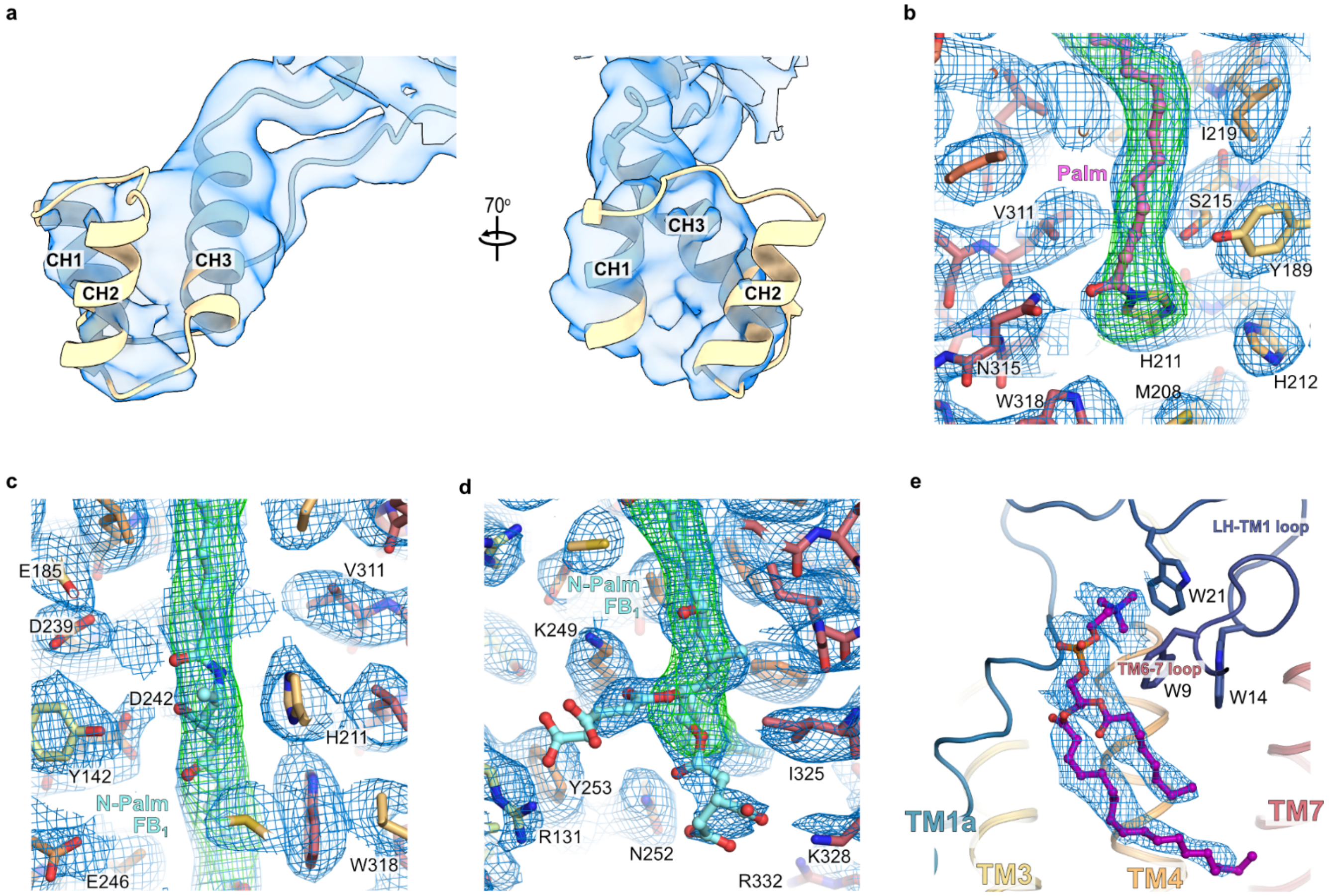


**Extended Data Figure 3. Cryo-EM map quality. a,** Fit of the Hox-like domain into the unsharpened cryo-EM map (blue surface). **b-d**, Cryo-EM density in the regions around the modelled ligands. The respective sharpened cryo-EM maps (blue mesh) and Servalcat Fo-Fc difference maps (obtained by omitting the acyl-imidazole and the N-palmitoyl fumonisin $B_1$ species from the models; green mesh) are overlaid on the models. **e,** Cryo-EM density for the bound lipid molecule, modelled as phosphatidylcholine (purple carbon atoms).

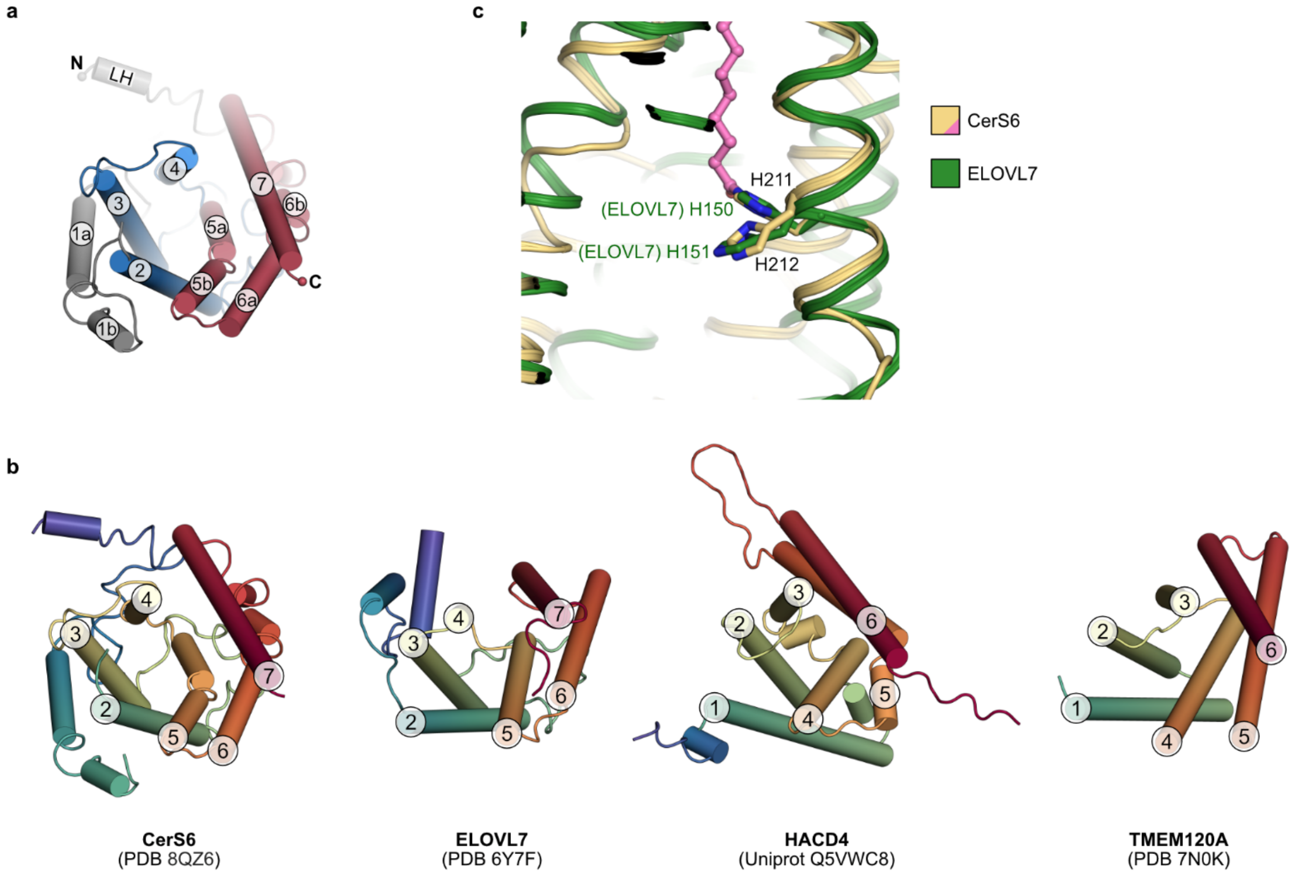


**Extended Data Figure 4. Transmembrane helix topology of CerS6. a**, The 6-TM barrel formed by TM2-7 of CerS6 is composed of two 3-TM units (TM2-4, blue; TM5-7, red), arranged as inverted repeats. **b**, Comparison of the transmembrane helix topology of the 6-TM barrels of CerS6, ELOVL7, HACD4 and TMEM120A (TACAN). **c,** Structural alignment of CerS6 and ELOVL7 reveals that their histidine pairs are structurally homologous. The acyl chain linked to His211 in the CerS6 covalent intermediate structure is shown in pink.

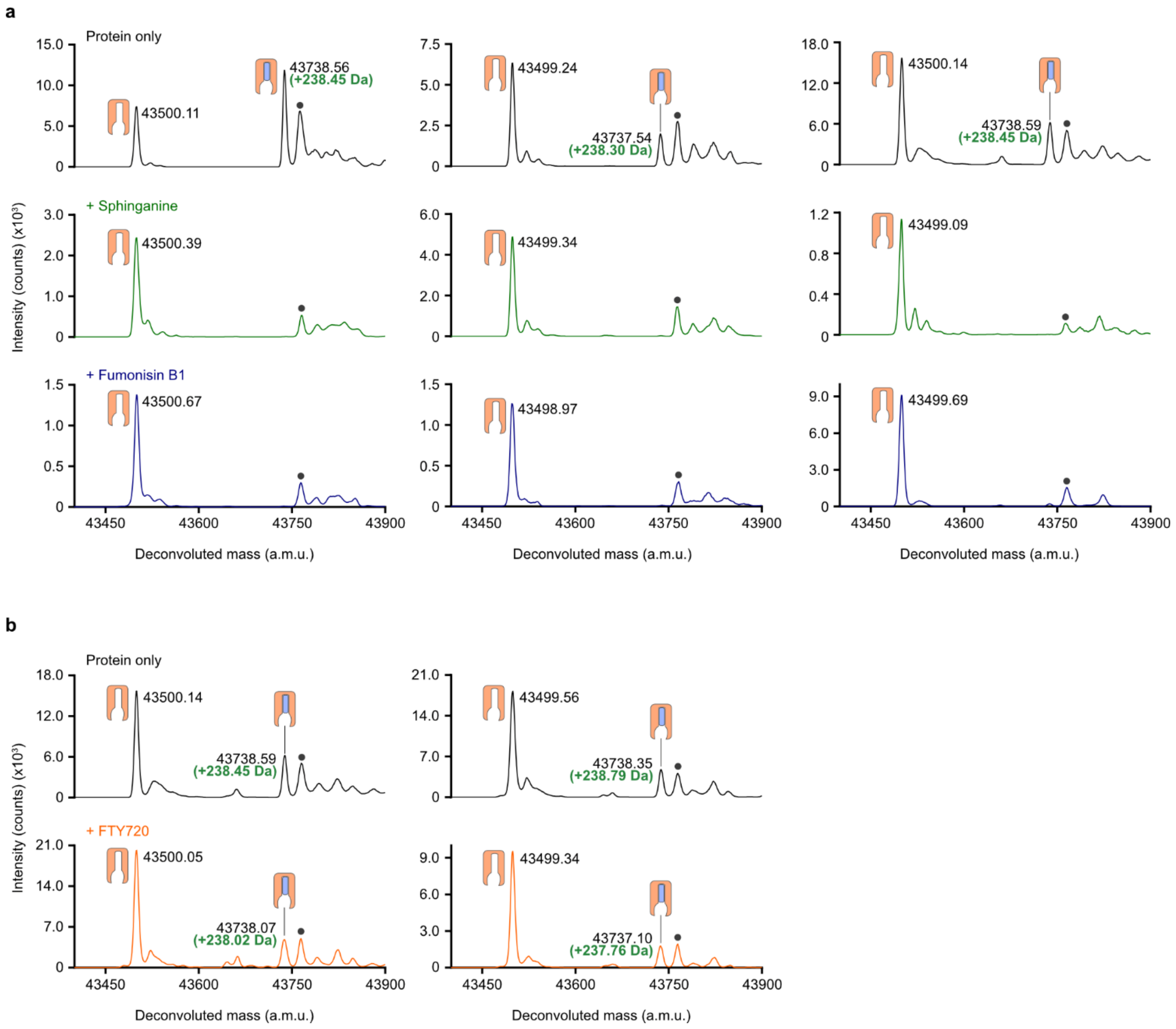


**Extended Data Figure 5. Monitoring the reaction of the covalent acyl-enzyme intermediate by intact protein MS.** Purified CerS6 was incubated in the presence and absence of substrates prior to liquid chromatography – electrospray ionization – mass spectrometry (LC-ESI-MS) intact mass analysis. **a,** Deconvoluted mass spectra are shown for CerS6 incubated in the absence of substrates (protein only), with 200 µM sphinganine, or with 200 µM fumonisin $B_1$. Deconvoluted mass peaks are indicated as follows: unmodified enzyme, orange icon; covalent acyl-enzyme species, orange and pink icon; background species (+264 Da) present in all traces, gray circle. Spectra are shown for 3 biological replicates. **b,** Deconvoluted mass spectra shown for CerS6 incubated in the presence and absence of 600 µM FTY720, revealing that the covalent acyl-enzyme species was preserved upon incubation with FTY720. Spectra are shown for 2 biological replicates.

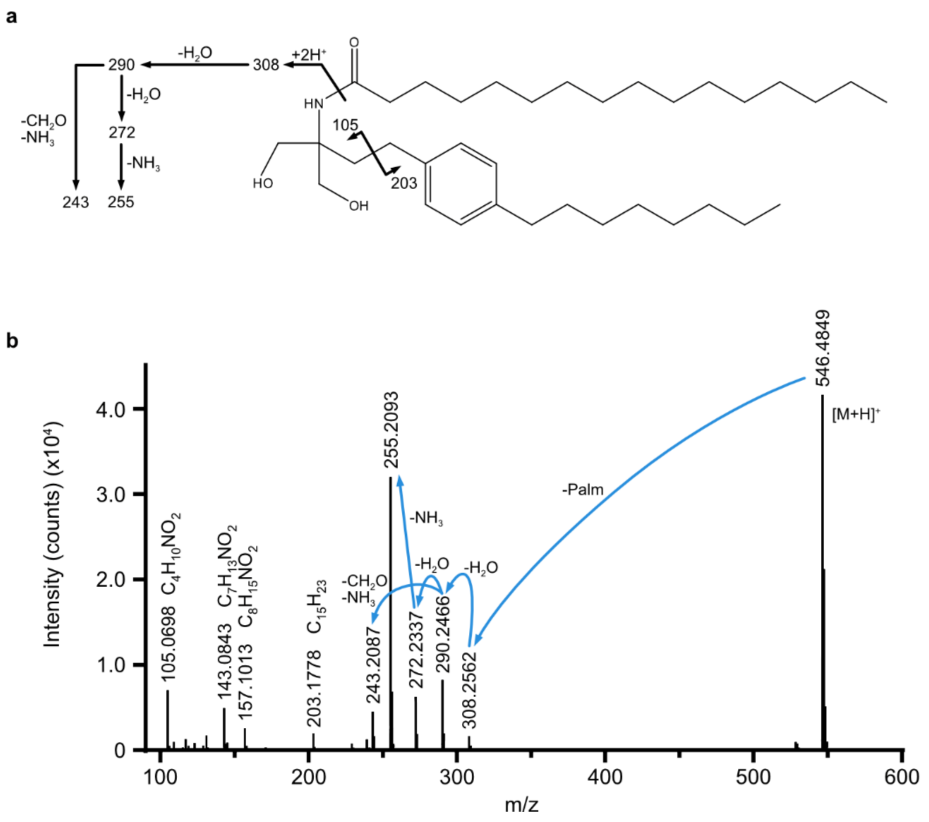


**Extended Data Figure 6. Structural characterization of N-palmitoyl FTY720 species by LC-ESI-MS/MS. a**, Structure of the proposed N-palmitoyl FTY720 reaction product, annotated with the m/z values of the daughter ions observed in panel b. **b,** MS/MS spectrum obtained from the fragmentation of the $[M+H]^+$ ion of N-palmitoyl FTY720 (m/z 546.4881) using a collision-induced dissociation energy of 14 V. Proposed relationships between the observed m/z peaks are indicated.

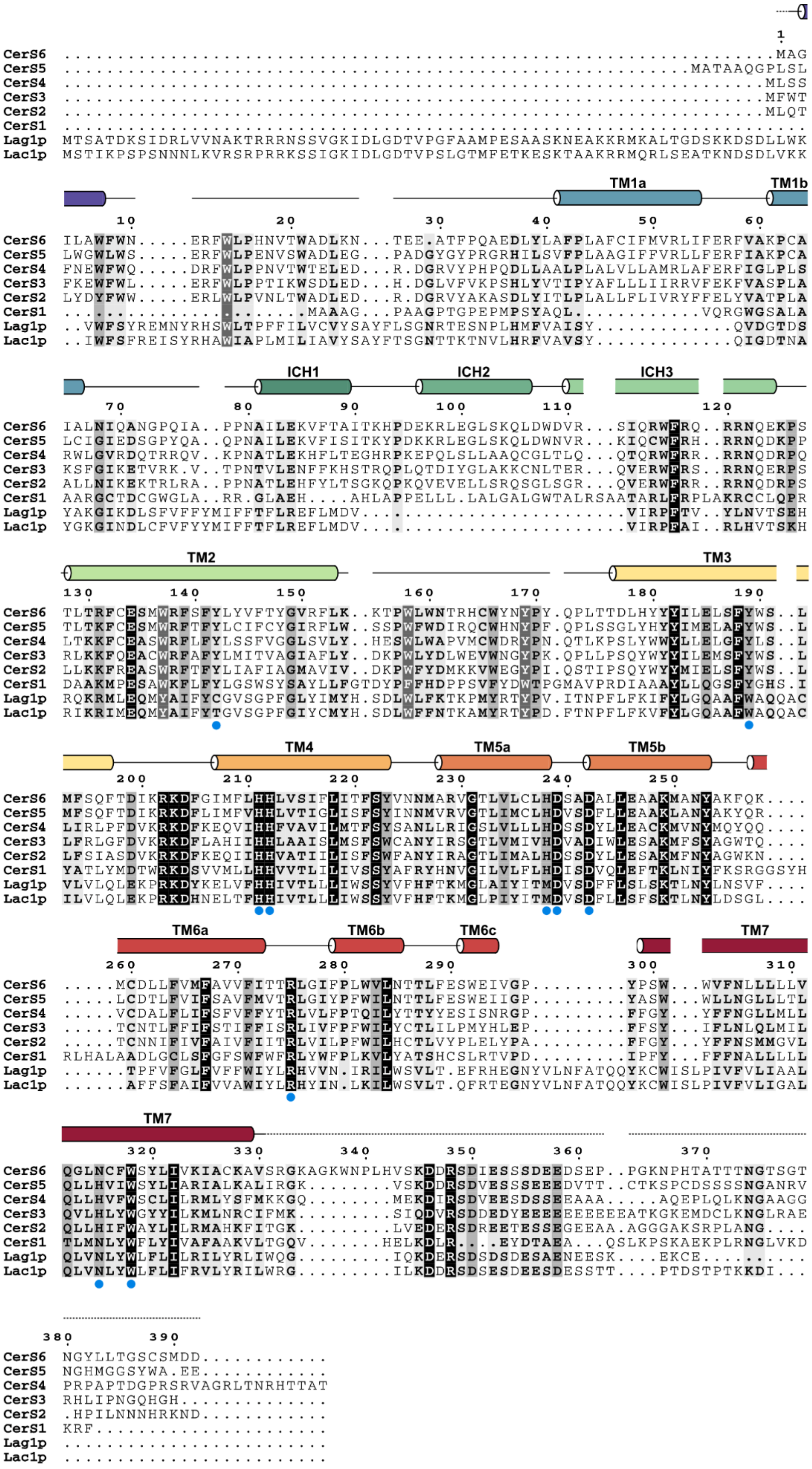


**Extended Data Figure 7. Multiple sequence alignment of human (CerS1-6) and *S. cerevisiae* (Lag1p, Lac1p) ceramide synthases.** Blue circles below the alignment indicate the active site residues.

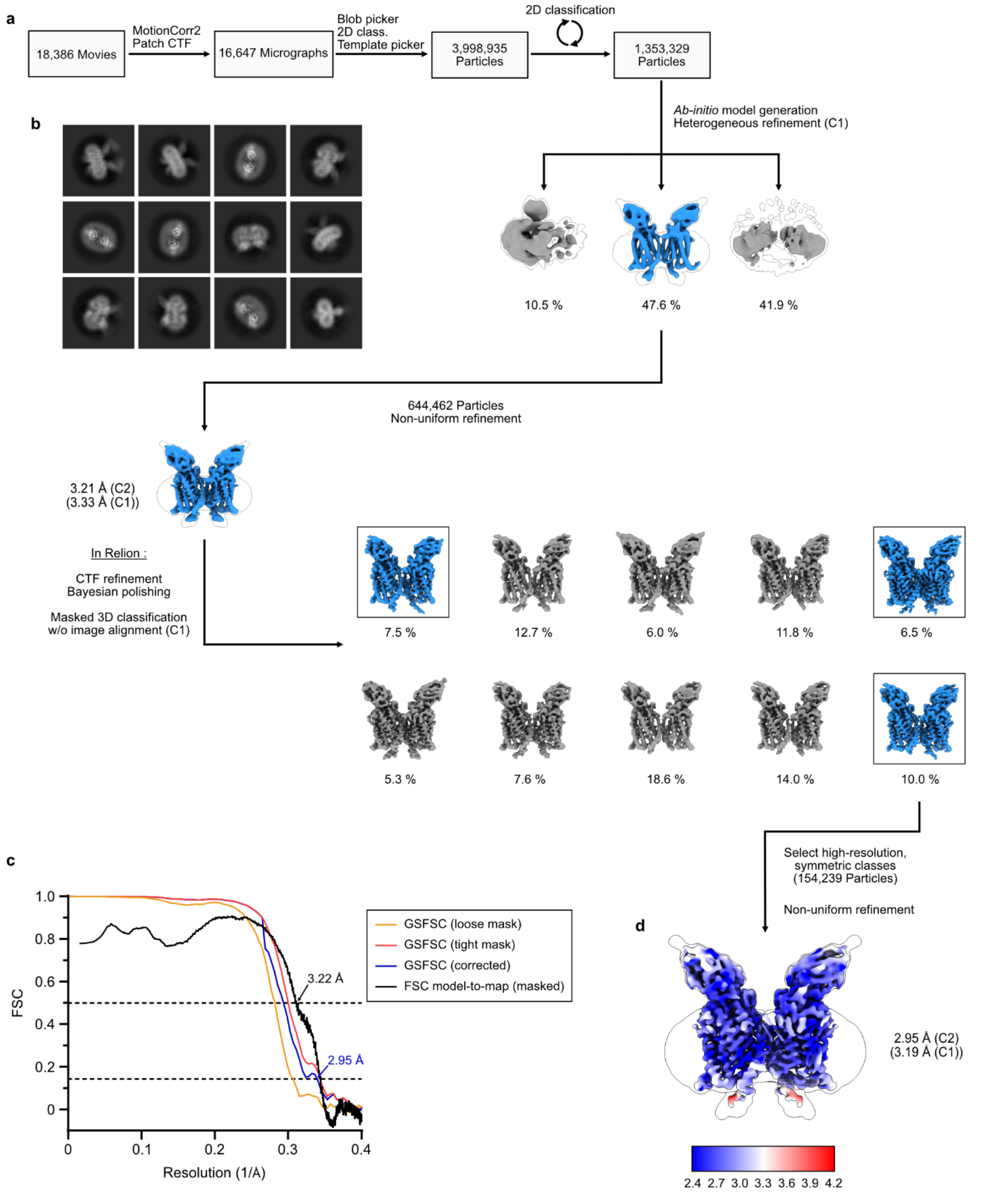


**Extended Data Figure 8. Cryo-EM data processing: N-palmitoyl FB$_1$-bound dataset**. **a**, Cryo-EM data processing flowchart. The final 3D reconstruction is colored by local resolution. **b**, 2D classes obtained prior to 3D classification. **c**, Fourier Shell Correlation (FSC) plots, indicating overall map resolution (GSFSC = 0.143) and a model-to-map FSC curve. **d,** Final 3D reconstruction, colored by local resolution.

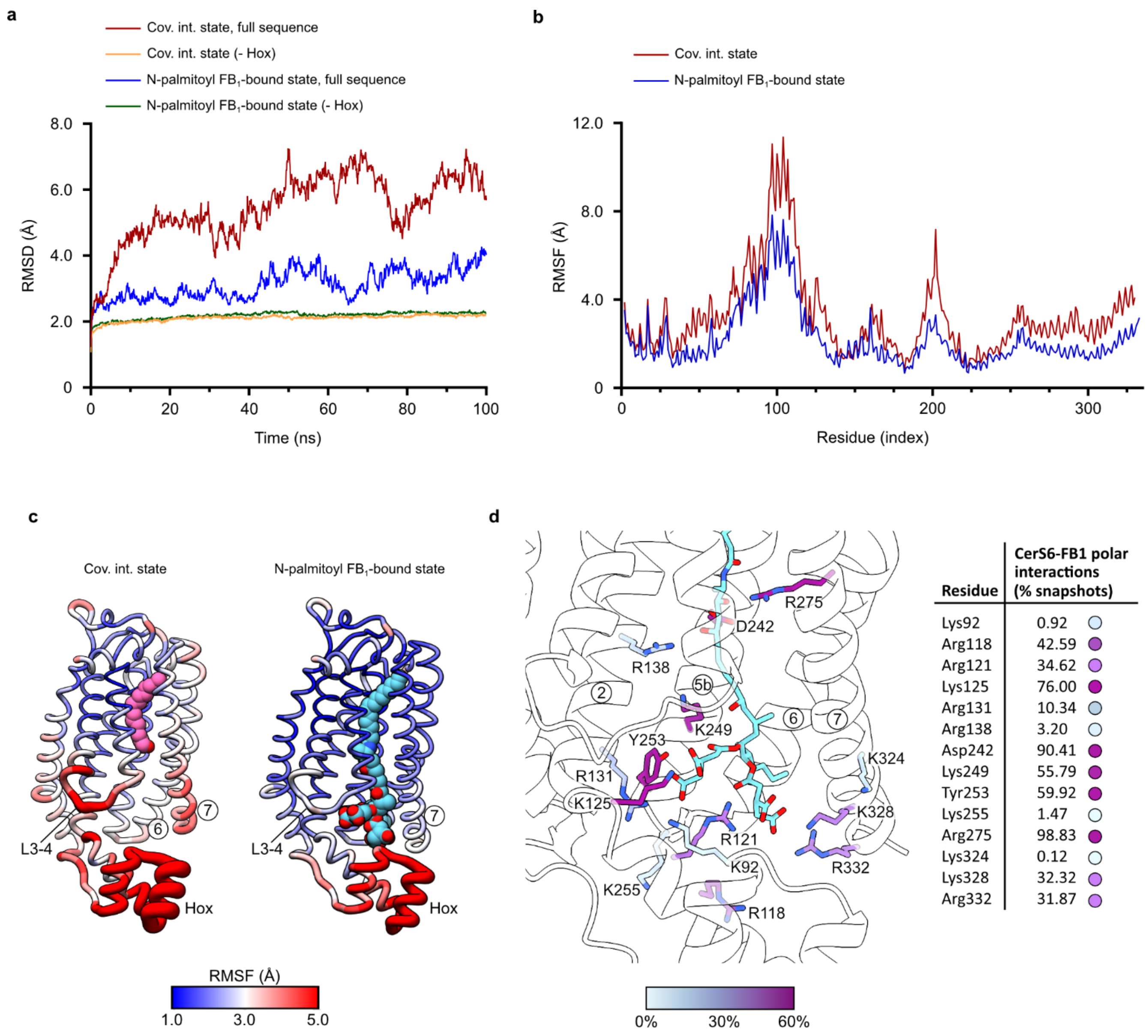


**Extended Data Figure 9. CerS6 dynamics and N-palmitoyl fumonisin $B_1$ interactions in simulations. a,** Average root-mean-square deviation (RMSD) values of Cα atoms from atomistic simulations of the covalent intermediate and N-palmitoyl fumonisin $B_1$-bound states. Traces where the Hox-like domain residues (70-126) were excluded from the RMSD calculation are shown to highlight the stability of the transmembrane region. **b,** Root-mean-square fluctuation (RMSF) values by residue (Cα atoms). RMSD and RMSF values shown are averaged over 4 x 100 ns atomistic simulations. **c,** N-palmitoyl $FB_1$ stabilises the CerS6 structure. RMSF values from the simulations are mapped onto the structures, revealing a reduction in the overall dynamics of the protein upon binding the reaction product N-palmitoyl $FB_1$, particularly in the Hox-like domain, the TM3-4 loop, TM6, and TM7. **d,** Analysis of the polar interactions between CerS6 and $FB_1$ during the simulations. Values shown correspond to the percentage of snapshots (taken every 100 ps during the simulations)

containing CerS6-$FB_1$ polar interactions at each residue. Residues are coloured by the frequency of polar interactions with $FB_1$ (cyan sticks).

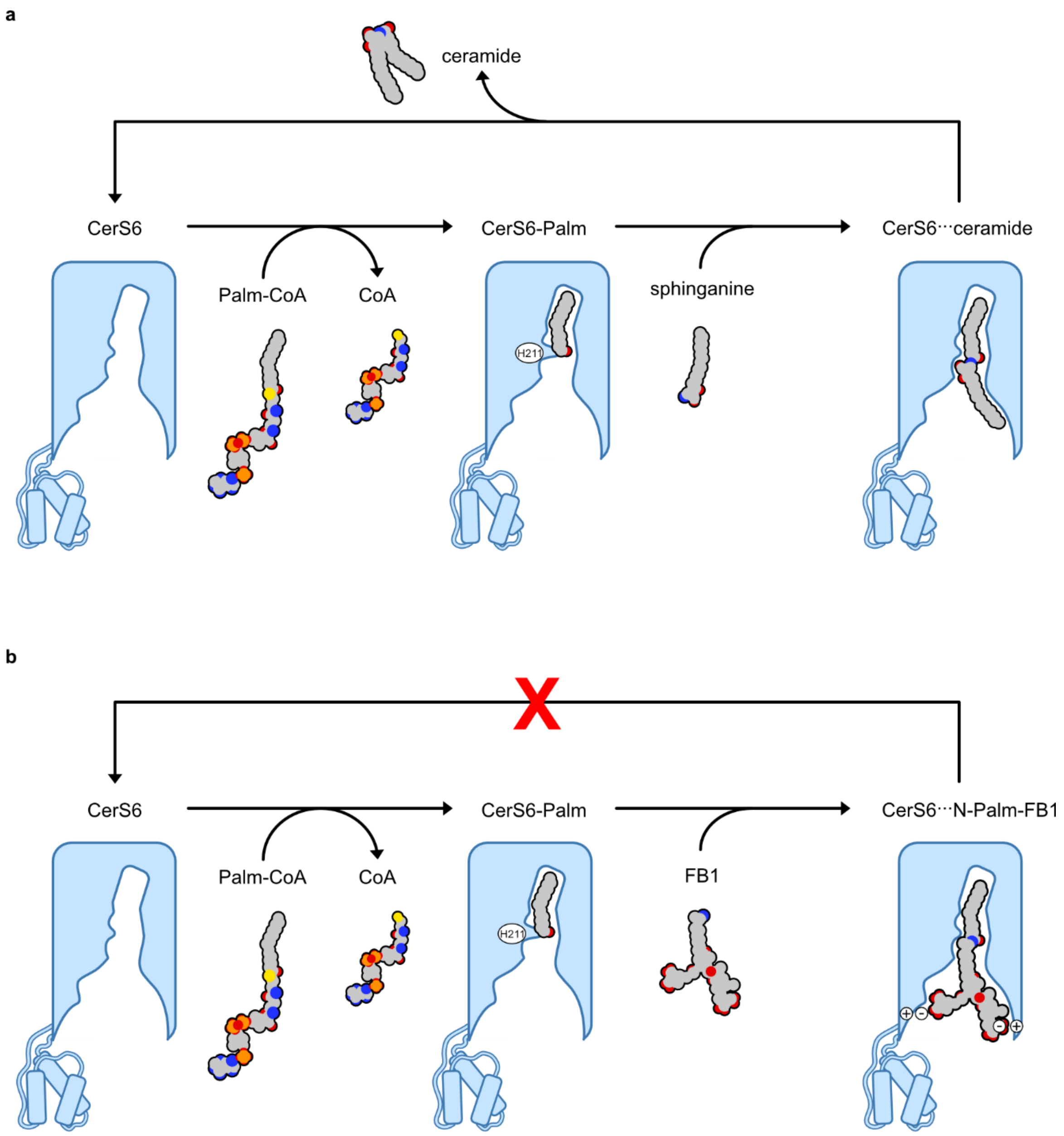


**Extended Data Figure 10. Cartoon representation of the proposed mechanism of ceramide synthases and their inhibition by FB$_1$. a,** CerS6 initially reacts with an acyl-CoA substrate, forming a covalent acyl-enzyme intermediate and releasing CoA as the first product. Subsequently, the reaction of the acyl-enzyme intermediate with sphinganine produces ceramide as the final product. Ceramide dissociates from the enzyme, which can then react with another acyl-CoA substrate to re-start the cycle. **b,** Reaction of the covalent acyl-enzyme intermediate species with FB$_1$ results in the formation of an N-acyl FB$_1$ product. This inhibitory product remains tightly bound in the central cavity, forming polar interactions near the cytoplasmic entrance, and preventing recycling of the enzyme.
